# The dual function monoclonal antibodies VIR-7831 and VIR-7832 demonstrate potent in vitro and in vivo activity against SARS-CoV-2

**DOI:** 10.1101/2021.03.09.434607

**Authors:** Andrea L. Cathcart, Colin Havenar-Daughton, Florian A. Lempp, Daphne Ma, Michael A. Schmid, Maria L. Agostini, Barbara Guarino, Julia Di iulio, Laura E. Rosen, Heather Tucker, Joshua Dillen, Sambhavi Subramanian, Barbara Sloan, Siro Bianchi, Dora Pinto, Christian Saliba, Katja Culap, Jason A Wojcechowskyj, Julia Noack, Jiayi Zhou, Hannah Kaiser, Sooyoung Lee, Nisar Farhat, Arthur Chase, Martin Montiel-Ruiz, Exequiel Dellota, Arnold Park, Roberto Spreafico, Anna Sahakyan, Elvin J. Lauron, Nadine Czudnochowski, Elisabetta Cameroni, Sarah Ledoux, Yoshihiro Kawaoka, Adam Werts, Christophe Colas, Leah Soriaga, Amalio Telenti, Lisa A. Purcell, Seungmin Hwang, Gyorgy Snell, Herbert W. Virgin, Davide Corti, Christy M. Hebner

## Abstract

Sotrovimab (VIR-7831) and VIR-7832 are dual action monoclonal antibodies (mAbs) targeting the spike glycoprotein of severe acute respiratory syndrome coronavirus 2 (SARS-CoV-2). Sotrovimab and VIR-7832 were derived from a parent antibody (S309) isolated from memory B cells of a 2003 severe acute respiratory syndrome coronavirus (SARS-CoV) survivor. Both mAbs contain an “LS” mutation in the Fc region to prolong serum half-life. In addition, VIR-7832 encodes an Fc GAALIE mutation that has been shown previously to evoke CD8+ T-cells in the context of an in vivo viral respiratory infection. Sotrovimab and VIR-7832 neutralize wild-type and variant pseudotyped viruses and authentic virus in vitro. In addition, they retain activity against monoclonal antibody resistance mutations conferring reduced susceptibility to previously authorized mAbs. The sotrovimab/VIR-7832 epitope continues to be highly conserved among circulating sequences consistent with the high barrier to resistance observed in vitro. Furthermore, both mAbs can recruit effector mechanisms in vitro that may contribute to clinical efficacy via elimination of infected host cells. In vitro studies with these mAbs demonstrated no enhancement of infection. In a Syrian Golden hamster proof-of concept wildtype SARS-CoV-2 infection model, animals treated with sotrovimab had less weight loss, and significantly decreased total viral load and infectious virus levels in the lung compared to a control mAb. Taken together, these data indicate that sotrovimab and VIR-7832 are key agents in the fight against COVID-19.

## INTRODUCTION

The coronavirus disease (COVID-19) pandemic caused by severe acute respiratory syndrome coronavirus 2 (SARS-CoV-2) has resulted in nearly 500 million confirmed cases and over 6.1 million deaths worldwide^1^. SARS-CoV-2 infection results in a broad range of disease severity^2^ Infection fatality rates increase significantly with age, with 28.3% of COVID-19 patients over the age of 85 succumbing to disease^2^. However, even in mild-to-moderate COVID-19 patients, significant post-infection sequelae can affect overall health and cause long-term disability^3^. While multiple SARS-CoV-2 vaccines are available for use, issues of supply, vaccine hesitancy and emergence of variants have been barriers to immunity globally^4–9^. In addition, there are individuals who remain at risk despite vaccination due to disease or underlying immunodeficiency. Thus, additional interventions and potential prophylactic agents continue to be needed to reduce morbidity and mortality due to COVID-19.

Over the course of the pandemic, multiple monoclonal antibodies (mAbs) targeting the SARS-CoV-2 spike protein have been authorized or approved for use in early treatment of COVID-19 patients^10–13^. However, many of these agents are ineffective against the currently circulating Omicron variants, especially those antibodies that target the receptor binding motif (RBM) of the viral spike (S) glycoprotein^14–16^. Antibodies targeting highly conserved epitopes outside of the RBM may potentially better retain activity against ever-emerging SARS-CoV-2 variants long-term. Furthermore, in addition to viral neutralization, antibodies possessing potent effector function to aid in the killing of virally infected cells and the elicitation of T cell immunity could additionally assist in halting disease progression^17–19^.

Sotrovimab and VIR-7832 are dual action mAbs derived from the parent antibody S309 identified from a 2003 SARS-CoV survivor^20^. These mAbs target an epitope containing a glycan (at position N343) that is highly conserved within the Sarbecovirus subgenus in a region of the S receptor binding domain (RBD) that does not compete with angiotensin converting enzyme 2 (ACE2) binding^21^. The variable region of sotrovimab and VIR-7832 have been engineered for enhanced developability. In addition, both antibodies possess an Fc “LS” mutation that confers extended half-life by binding to the neonatal Fc receptor^22–24^. VIR-7832 is identical to sotrovimab with the exception of the addition of a 3 amino acid GAALIE (G236A, A330L, I332E) modification to the Fc domain^25^. The GAALIE modification has previously been shown in vitro to enhance binding to FcγIIa and FcγIIIa receptors, decrease affinity for FcγIIb compared to typical IgG1 and evoke protective CD8+ T-cells in the context of viral respiratory infection in vivo^26,27^.

Here we characterize the antiviral activity of sotrovimab and VIR-7832. These mAbs effectively neutralize SARS-CoV-2 live virus in vitro as well as in pseudotyped virus assays against emerging variants of concern and variants that confer resistance to previously authorized or approved mAbs^28^. In addition to the neutralizing capacity, both antibodies demonstrate potent effector function and mediate antibody dependent cellular cytotoxicity (ADCC) and antibody dependent cellular phagocytosis (ADCP) in vitro. Furthermore, resistance selection experiments and epitope conservation analyses indicate the potential for a high barrier to resistance. Data derived from the Syrian golden hamster model demonstrates efficacy in a proof-of-concept in vivo model. Taken together, these data indicate that sotrovimab and VIR-7832 are key components of the arsenal in the fight against COVID-19.

## RESULTS

### Sotrovimab and VIR-7832 bind SARS-CoV-2 spike and effectively neutralize live virus in vitro

Previously published work showed that S309 bound SARS-CoV-2 recombinant and cell surface-associated S and neutralized live virus in vitro^20^. We initiated these studies by repeating and extending these earlier results. To determine the binding activity of sotrovimab and VIR-7832 to the SARS-CoV-2 S, enzyme-linked immunosorbent assay (ELISA), surface plasmon resonance (SPR) and flow cytometry assays were utilized. Sotrovimab and VIR-7832 bound to recombinant S RBD (amino acids 331-541) with EC_50_ values of 20.40 ng/mL and 14.9 ng/mL, respectively, by ELISA (Figure 1a). Using SPR, both antibodies demonstrated potent binding to recombinant S RBD with an equilibrium constant (Kd) of 0.21 nM (Figure 1b). As antibody recognition of cell surface-bound S could mediate killing of virally infected cells, flow cytometry-based studies using cells transiently transfected with a S-encoding plasmid were used to examine antibody binding to cell surface-expressed S trimer. By this method, both sotrovimab and VIR-7832 bound efficiently to surface-expressed S (Figure 1c).

**Figure 1.**
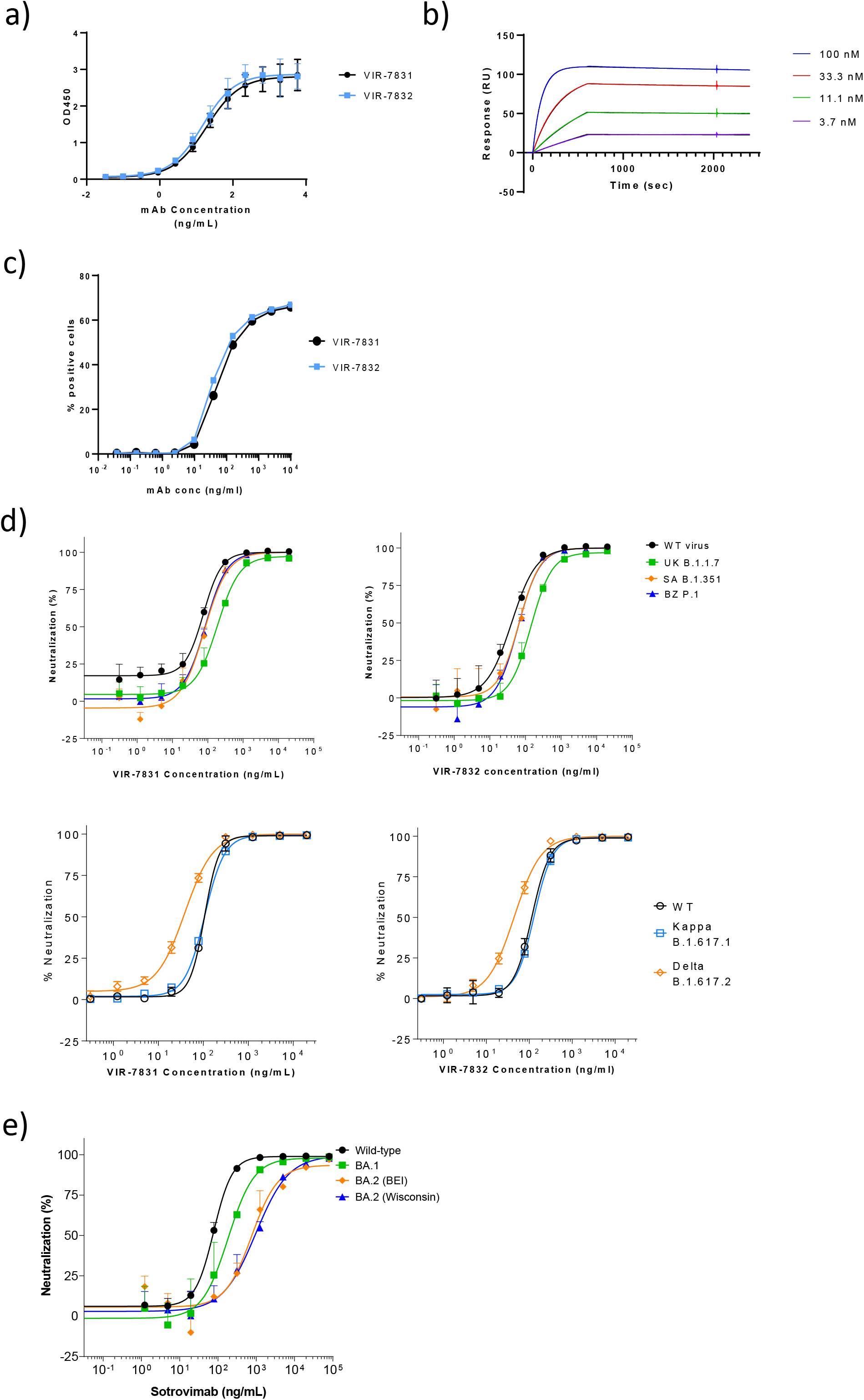
Sotrovimab (VIR-7831) and VIR-7831 bind S and neutralize SARS-CoV-2 virus and S variants in vitro. a) Binding of sotrovimab (VIR-7831; black circles) and VIR-7832 (blue squares) to SARS-CoV-2 RBD was tested by ELISA. Shown is the average of four replicates and SD derived from three independent experiments. b) Association and dissociation profiles of sotrovimab (VIR-7831) to SARS-CoV-2-RBD were measured using SPR. The double reference subtracted curves (shown for single replicates) are plotted together with the curve fit in black (obscured by close overlay with the data). Values are from two independent experiments. c) Binding of sotrovimab (VIR-7831; black circles) and VIR-7832 (blue squares) to cell-surface S protein was determined by flow cytometry. Data are expressed as the percentage of the positive cells. Results shown are from one experiment and representative of three independent experiments performed. d) In vitro neutralization of live SARS-CoV-2 by different concentrations of Sotrovimab (VIR-7831; left) and VIR-7832 (right) measured by nucleocapsid staining 6-hours post-infection. Results shown are from one experiment and representative of at least two independent experiments performed. e) In vitro neutralization of Omicron BA.2 authentic virus with BA.1 and wild-type (Washington) virus run as controls. Two unique Omicron BA.2 isolates originating from different sources were tested (BEI and University of Wisconsin). Results shown are from one experiment and are representative of at least three independent experiments performed.

To examine neutralization capacity, sotrovimab and VIR-7832 were tested in a VeroE6 or Vero E6-TMPRSS2 cell-based live SARS-CoV-2 virus system against the Washington 2019 (wild-type) virus as well as against the Alpha (B.1.1.7), Beta (B.1.351), Gamma (P.1), Delta (B.1.617.2), Kappa (B.1.617.1) and Omicron (BA.1, BA.1.1 and BA.2) variants. Concentration-dependent viral neutralization of the Washington 2019 strain was observed for both antibodies, with geometric mean IC_50/90_ values of 100.1/186.3 ng/mL and 78.3/253.1 ng/mL, respectively (Figure 1d). IC_50/90_ values observed for sotrovimab and VIR-7832 against the Beta, Gamma, Delta and Kappa variant viruses were similar to those against the wild-type strain. A slight shift in the sotrovimab/VIR-7832 IC_50/90_ compared to wild-type was observed for the Alpha and Omicron BA.1 and BA.1.1 variants (Figure 1d, 1e; Table 1). VIR-7832 had a 3.1-fold shift in IC_50_ and 3.7-fold shift in IC_90_ versus wild-type against the Alpha variant and was not tested against the Omicron variants. A shift in in vitro activity was observed for sotrovimab against two differentially sourced Omicron BA.2 variant isolates (BEI and University of Wisconsin) resulting in an average 15.7-fold and 35.1-fold shift in IC_50_ and IC_90_, respectively, compared to wild-type.

**Table 1.** Sotrovimab and VIR-7832 neutralize S variants of concern in an authentic virus system. Average fold change in sotrovimab and VIR-7832 IC_50_ compared to relative wild-type controls for S variants tested in an authentic virus system. Data shown are averages of at least two independent experiments. NT = not tested

| SARS-CoV-2 Variant | Geometric Mean Sotrovimab IC<sub>50</sub> (ng/ml)<br>(Average Fold Change IC<sub>50</sub> vs. Wild-Type) | Geometric Mean Sotrovimab IC<sub>90</sub> (ng/ml)<br>(Average Fold Change IC<sub>90</sub> vs. Wild-Type) | Geometric Mean VIR-7832 IC<sub>50</sub> (ng/ml)<br>(Average Fold Change IC<sub>50</sub> vs. Wild-Type) | Geometric Mean VIR-7832 IC<sub>90</sub> (ng/ml)<br>(Average Fold Change IC<sub>90</sub> vs. Wild-Type) |
| --- | --- | --- | --- | --- |
| Alpha (B.1.1.7) | 187.2 (3.0) | 1246.9 (4.1) | 181.1 (3.1) | 1223.0 (3.7) |
| Beta (B.1.351) | 71.9 (1.2) | 385.0 (1.3) | 59.67 (1.1) | 309.84 (0.9) |
| Gamma (P.1) | 73.11 (1.6) | 335.79 (1.4) | 48.9 (1.2) | 217.0 (0.9) |
| Delta (B.1.617.2) | 51.3 (0.4) | 219.1 (0.6) | 59.13 (0.5) | 274.24 (0.7) |
| Kappa (B.1.617.1) | 118.64 (0.9) | 379.89 (1.0) | 132.4 (1.0) | 428.8 (1.0) |
| Omicron (BA.1) | 169.2 (3.8) | 944.1 (4.6) | NT | NT |
| Omicron (BA.1.1) | 165.1 (4.3) | 827.1 (4.2) | NT | NT |
| Omicron (BA.2) | 972.8 (15.7) | 9476.3 (35.1) | NT | NT |

As variant evolution is a natural part of SARS-CoV-2 biology and emerging live virus variants are not always readily accessible for testing, a vesicular stomatitis virus (VSV)-based pseudotyped virus system targeting Vero E6 cells was used to examine sotrovimab and VIR-7832 neutralization against emergent variants (Table 2). Fold-changes in sotrovimab and VIR-7832 IC_50_ values against pseudotyped virus expressing spike from the Alpha, Beta, Gamma, Delta, Kappa and Omicron variants were similar to those observed in the authentic virus system. Sotrovimab was tested against an extended panel of pseudotyped viruses incorporating emerging variants as well as variants currently or previously deemed as Variants of Concern (VOC) or Variants of Interest (VOI) by the World Health Organization (WHO). Sotrovimab neutralized all variants tested with fold changes in IC_50_s of ≤3.3-fold observed for 21 of 23 variants assessed (Table 2), while a 7.3-fold change in IC_50_ was observed for Omicron BA.3 and a 16-fold change in IC_50_ was observed for the Omicron BA.2 variant (Table 2).

**Table 2.** Sotrovimab and VIR-7832 retain activity against S variants of concern in a pseudotyped virus system. Average fold change in sotrovimab and VIR-7832 IC_50_ compared to relative wild-type controls for S variants tested in a VSV/VeroE6 pseudotyped virus system. Data shown are averages of at least two independent experiments.

| Variant | Spike Mutations | Fold-Change in Sotrovimab IC<sub>50</sub> vs. Wild-type | Fold-Change in VIR-7832 IC<sub>50</sub> vs. Wild-type |
| --- | --- | --- | --- |
| Alpha (B.1.1.7) | H69-, V70-, Y144-, N501Y, A570D, D614G, P681H, T716I, S982A, D1118H | 2.3 | 2.5 |
| Beta (B.1.351) | L18F, D80A, D215G, R246I, K417N, E484K, N501Y, D614G, A701V | 0.6 | 0.7 |
| Gamma (P.1) | D138Y, D614G, E484K, H655Y, K417T, L18F, N501Y, P26S, R190S, T1027I, T20N, V1176F | 0.4 | 0.4 |
| Delta (B.1.617.2) | T19R, G142D, E156G, F157-, R158-, L452R, T478K, D614G, P681R, D950N | 1 | NT |
| Epsilon (B.1.427/B.1.429) | S13I, W152C, L452R, D614G | 0.7 | NT |
| Eta (B.1.525) | Q52R, A67V, H69-, V70-, Y144-, E484K, D614G, Q677H, F888L | 0.9 | NT |
| Iota (B.1.526) | L5F, T95I, D253G, E484K, D614G, A701V | 0.6 | NT |
| Kappa (B.1.617.1) | T95I, G142D, E154K, L452R, E484Q, D614G, P681R, Q1071H | 0.7 | NT |
| Lambda (C.37) | G75V, T76I, del246-252, L452Q, F490S, T859N | 1.5 | NT |
| Mu (B.1.621) | T95I, Y144T, Y145S, ins146N, R346K, E484K, N501Y, D614G, P681H, D950N | 1.3 | NT |
| Omicron (BA.1) | A67V, H69del, V70del, T95I, G142D, V143del, Y144del, Y145del, N211del, L212I, ins214EPE, G339D, S371L, S373P, S375F, K417N, N440K, G446S, S477N, T478K, E484A, Q493R, G496S, Q498R, N501Y, Y505H, T547K, D614G, H655Y, N679K, P681H, N764K, D796Y, N856K, Q954H, N969K, L981F | 2.7 | NT |
| Omicron + R346K (BA.1.1) | A67V, H69del, V70del, T95I, G142D, V143del, Y144del, Y145del, N211del, L212I, ins214EPE, G339D, R346K, S371L, S373P, S375F, K417N, N440K, G446S, S477N, T478K, E484A, Q493R, G496S, Q498R, N501Y, Y505H, T547K, D614G, H655Y, N679K, P681H, N764K, D796Y, N856K, Q954H, N969K, L981F | 3.3 | NT |
| Omicron (BA.2) | T19I, del24-26, A27S, G142D, V213G, G339D, S371F, S373P, S375F, T376A, D405N, R408S, K417N, N440K, S477N, T478K, E484A, Q493R, Q498R, N501Y, Y505H, D614G, H655Y, N679K, P681H, N764K, D796Y, Q954H, N969K | 16.0 | NT |
| Omicron (BA.3) | A67V, del69-70, T95I, G142D, del143-145, del211, L212I, G339D, S371F, S373P, S375F, D405N, K417N, N440K, G446S, S477N, T478K, E484A, Q493R, Q498R, N501Y, Y505H, D614G, H655Y, N679K, P681H, N764K, D796Y, Q954H, N969K | 7.3 | NT |
| Delta Plus (AY.1) | T19R, T95I, G142D, E156G, F157-, R158-, W258L, K417N, L452R, T478K, D614G, P681R, D950N | 1.1 | NT |
| Delta Plus (AY.2) | T19R, V70F, G142D, E156G, F157-, R158-, A222V, K417N, L452R, T478K, D614G, P681R, D950N | 1.3 | NT |
| Delta Plus (AY. 4.2) | T19R, T95I, G142D, Y145H, E156G, F157, R158-, A222V, L452R, T478K, D614G, P681R, D950N | 1.6 | NT |
| Mexico/Swiss (B.1.1.519) | T478K, D614G, P681H, T732A | 0.8 | NT |
| Scotland (B.1.258) | H69-, V70-, N439K, D614G | 0.9 | NT |
| US (R.2) | E484K, D614G, Q677H, T732S, E1202Q | 0.8 | NT |
| Liverpool (A.23.1) | R102I, F157L, V367F, E484K, Q613H, P681R | 1.1 | NT |
| Cameroon (B.1.619) | I210T, N440K, E484K, D614G, D936N, S939F, T1027I | 1.3 | NT |
| Bristol (B.1.1.7+E484K) | H69-, V70-, Y144-, E484K, N501Y, A570D, D614G, P681H, T716I, S982A, D1118H | 1.7 | NT |

### Sotrovimab and VIR-7832 exhibit potent effector function in vitro

Although direct antiviral mechanisms are crucial to provide protection, Fc-dependent mechanisms mediated by interaction with Fc gamma receptors (FcγRs) on immune cells or with complement, can contribute to overall potency in vivo. The potential for sotrovimab and VIR-7832 to mediate effector functions were assessed in vitro by measuring binding to FcγRs and C1q and in assays designed to demonstrate antibody-dependent cellular cytotoxicity (ADCC) or antibody-dependent cellular phagocytosis (ADCP)^29–32^.

Antibody binding to the human activating FcγRIIa (low-affinity R131 and high affinity H131 alleles), FcγRIIIa (low-affinity F158 and high-affinity V158 alleles), and to the inhibitory FcγRIIb were examined using SPR (Supplemental figure 1a). Sotrovimab similarly bound both the H131 and R131 alleles of FcγRIIa and binds FcγRIIb. Sotrovimab bound both FcγRIIIa alleles, with reduced binding to the F158 allele compared to V158, as is characteristic for human IgG1^33^. Binding of sotrovimab to C1q was similar to the parental antibody (S309-LS) (Supplemental figure 1b). As previously reported for antibodies encoding the GAALIE mutation^25, 34^, VIR-7832 bound with comparatively higher affinity to activating FcγRIIa and FcγRIIIa than sotrovimab (Supplemental figure 1a). Conversely, VIR-7832 showed reduced affinity for FcγIIb and abrogation of binding to C1q (Supplemental figure 1b).

The antibodies were also assessed for the ability to activate human FcγRIIa, FcγRIIb or FcγRIIIa, using a Jurkat cell reporter assay^35^ (Figure 2a-d). S309-GRLR, which contains the effector function-abrogating G236R, L328R mutations was used as a negative control. Cells stably transfected with the SARS-CoV-2 spike protein (CHO-CoV-2-Spike) served as target cells. Both sotrovimab and the parental S309-LS activated signaling of the higher-affinity allele FcγRIIa (H131) but did so less efficiently than the GAALIE-containing antibody VIR-7832 (Figure 2a) while sotrovimab, VIR-7832 and S309-LS induced similar low-level activation of the inhibitory receptor FcγRIIb (Figure 2b). Sotrovimab demonstrated substantially lower activation of FcγRIIIa F158 versus V158 as expected while VIR-7832 showed increased activation of both alleles of FcγRIIIa (F158 and V158) (Figures 2c,d).

**Figure 2.**
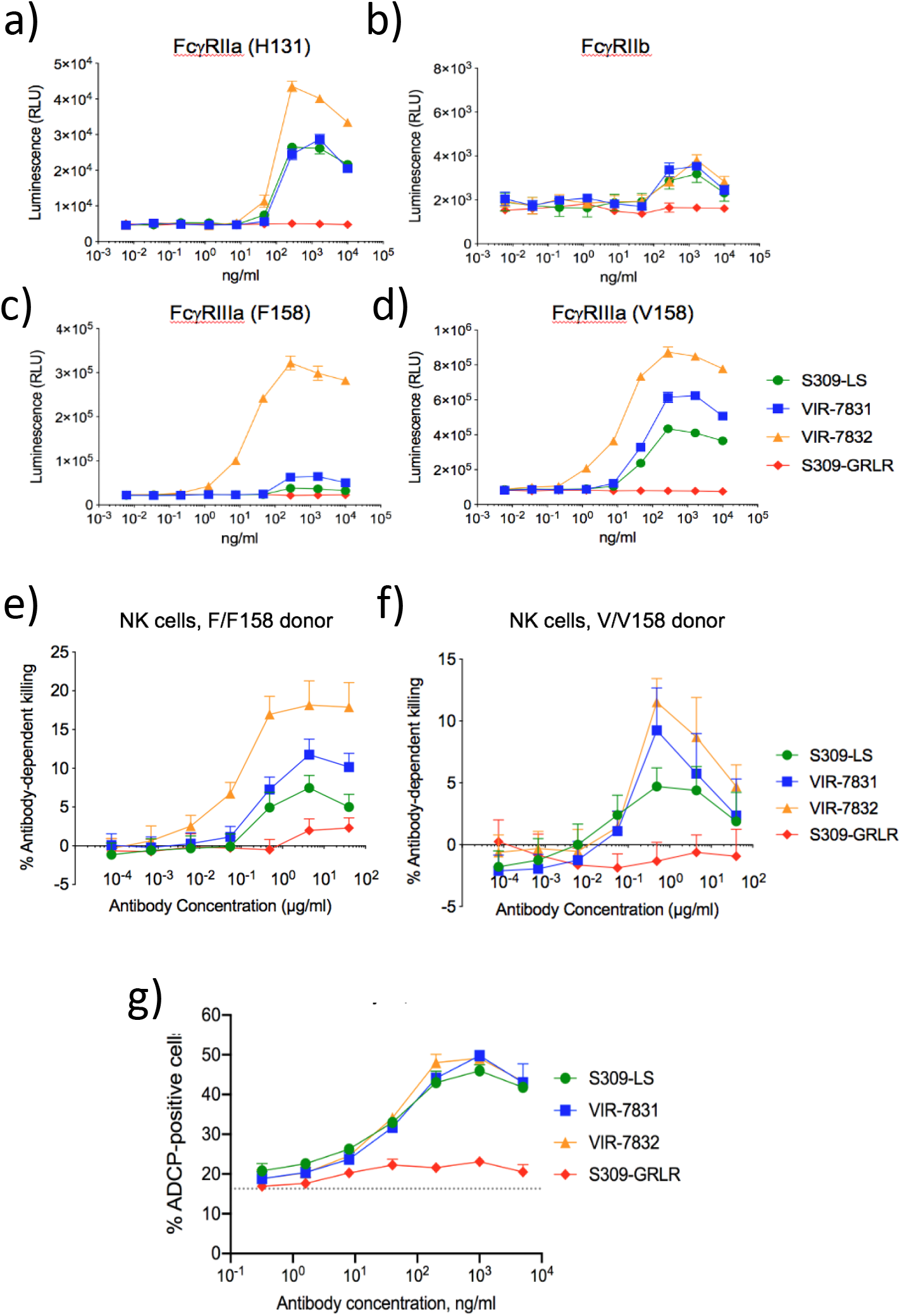
Sotrovimab (VIR-7831) and VIR-7832 demonstrate effector function in vitro. In vitro effector function (a-e) activation profiles of human FcγRIIa (a), FcγRIIb (b), FcγRIIIa low-affinity (F158) (c) or FcγRIIIa high-affinity binding allele (V158) (d) using bioreporter assays using S-expressing CHO cells as the target antigen. Data points show means± SD of duplicates. NK-cell mediated killing (ADCC) of S-expressing CHO cells using freshly isolated cells from two donors previously genotyped for homozygous expression of low-affinity (F/F158) (e) or high-affinity (V/V158) FcγRIIIa (f). Data points are means of quadruplicates ± SD. g) Antibody-dependent cellular phagocytosis (ADCP) using S-expressing CHO cells and freshly isolated PBMCs. Data represent the means of duplicates ± SD.

To further elucidate the effector function potential of the antibodies, ADCC and ADCP assays were performed using donor PBMCs or NK cells as effector cells and CHO cells stably expressing S (CHO-CoV-2-Spike) as target cells (Figure 2e-g). The ability of antibodies to activate NK cell-mediated killing was measured in vitro using two genotyped donors expressing homozygous low-affinity (F/F158) or high-affinity (V/V158) (Figure 2e-f). Compared to the parental mAb S309-LS, sotrovimab had slightly increased capacity to induce NK cell-mediated ADCC when using cells from either F/F158 or V/V158 donors. As expected, VIR-7832 induced NK cell-mediated ADCC in cells from donors expressing the low-affinity F/F158 allele of FcγIIIa more efficiently than VIR-7831. These results were confirmed with NK cells from a heterozygous donor (F/V158).

The ability of sotrovimab and VIR-7832 to facilitate ADCP by primary CD14^+^ monocytes was measured in vitro by exposing freshly isolated human PBMCs to CHO-CoV-2-Spike cells that were pre-incubated with antibody (Figure 2g). Sotrovimab, VIR-7832 and S309-LS induced similar levels of ADCP by CD14^+^ monocytes. These results indicate that sotrovimab and VIR-7832 have the potential to trigger ADCC and ADCP of cells displaying SARS CoV-2 S protein.

### Subneutralizing levels of sotrovimab and VIR-7832 do not enhance virus uptake, replication or cytokine production in vitro

One potential concern with any antibody therapeutic targeting a viral agent is the possibility of antibody-dependent enhancement (ADE). ADE is an in vivo phenomenon in which the presence of an antibody worsens disease. There are several in vitro assays that may provide plausible correlates for ADE in vivo, though none of these have been proven relevant to COVID-19 as to date ADE has not been observed in trials of monoclonal antibodies or plasma^13, 36–38^. ADE can occur by several potential mechanisms^39^. Poorly neutralizing antibodies or subneutralizing levels of antibody could theoretically facilitate enhanced virus entry and infection through Fc receptor interactions. A second theoretical mechanism involves antibody-antigen complex formation leading to enhanced cytokine production. A third mechanism of ADE has been observed in a porcine model of influenza where the kinetics of viral fusion to the target cell was enhanced in a Fab-dependent manner by fusion-enhancing non-neutralizing antibodies^40, 41^.

To explore whether sotrovimab and VIR-7832 exhibit in vitro activities that might be related to ADE in vivo, we evaluated SARS-CoV-2 replication in human cells that express FcγRs: monocyte-derived dendritic cells (moDCs), peripheral blood mononuclear cells (PBMCs) and the human U937 macrophage cell line (Supplemental Figure 2a-b). Subneutralizing concentrations of sotrovimab and VIR-7832 were precomplexed with SARS-CoV-2 (MOI =0.01) and added to target cells. Using immunostaining methods, at 24 hours post-infection no productive entry of SARS-CoV-2 into moDCs, PBMCs, or U937 cells was observed in the presence or absence of either mAb, while VeroE6 control cells demonstrated internalization in all conditions evaluated. Reduced internalization of SARS-CoV-2 in VeroE6 cells was observed at the highest concentration of sotrovimab and VIR-7832 (p-value <0.05), indicating effective virus neutralization prevented virus entry. Using a focus forming assay, virus replication and secretion of infectious virus were detectable by 48 hours post-infection in VeroE6 cells, with comparable levels of replication in the presence or absence of sotrovimab or VIR-7832. However, no replication of SARS-CoV-2 was detected in moDCs, PBMCs or U937 cells regardless of antibody treatment, indicating lack of productive SARS-CoV-2 infection of these cells, consistent with previously published data^42^.

To evaluate the potential for sotrovimab and VIR-7832 to enhance cytokine release upon SARS-CoV-2 infection in FcγR-expressing cells, cytokines and chemokines were measured in the supernatants from cells infected with SARS-CoV-2-in the presence of sotrovimab or VIR-7832 (Supplemental figure 2c). Levels of IFN-γ, IL-10, IL-6, IL 8, IP-10, MCP-1, and TNF-α in the supernatant were quantified by MSD at 24-and 48-hours post-infection. For all cell types evaluated, cytokine/chemokine production was similar between all antibody concentrations tested and the no antibody control at both 24- and 48-hours post-infection. Taken together, these in vitro data indicate that neither VIR-7831 nor VIR-7832 exhibit in vitro activities that have been proposed to possibly correlate with ADE in vivo.

### Sotrovimab and VIR-7832 have a high barrier to resistance in vitro and do not display cross-resistance with other SARS-CoV-2 mAbs

We next determined whether resistant variants could be elicited by serial passage of SARS-CoV-2 in the presence of VIR-7832. As sotrovimab and VIR-7832 differ only in the Fc region of the antibody, resistance selection experiments were conducted with VIR-7832 as a proxy for both antibodies. SARS-CoV-2 was subjected to 10 passages in the presence of VIR-7832 at fixed concentrations of ∼10x, 20x, 50x or 100x IC_50_ (1, 2, 5, or 10 μg/mL) in VeroE6 cells. No CPE was detected in wells passaged with antibody through 10 passages, while CPE was observed in the no antibody control in all passages. Similarly, no virus was detected by focus forming assay at any concentration of VIR-7832 through all 10 passages even at the lowest concentration tested.

As no viral breakthrough was observed in the fixed concentration resistance selection, a second method was employed wherein SARS-CoV-2 virus was passaged in sub-IC_50_ concentrations of antibody followed by subsequent passaging in the presence of increasing concentrations of mAb in an attempt to force resistance emergence (Supplemental Figure 3). Passaging was performed in duplicate wells to account for founder effects, and concentration increases for each well were based on CPE observations. Five sequential passages were conducted using increasing concentrations of VIR-7832 at 0.5, 1, 2, 5 and ∼10x IC_50_ (0.05, 0.1, 0.2, 0.5, 1 μg/mL; Supplemental Figure 3a), though no CPE was observed by passages 4 and 5 (0.5 and 1 μg/mL, respectively) indicating that variants originally selected at the lower concentrations were either unfit or susceptible to the higher concentrations of antibody. To further assess whether resistance mutations could be generated, selection was restarted using passage 3 virus generated with ∼2x IC_50_ (0.2 μg/mL) of VIR-7832 in duplicate wells at ∼2x and ∼5x IC_50_ (0.2, 0.5 μg/mL), generating two passage lineages (Supplemental Figure 3b-c).

Supernatants were evaluated for detectable virus at each passage by focus forming assay and cell supernatants from viral passages containing detectable virus were tested in SARS-CoV-2 neutralization assays to evaluate IC_50_ shifts as a marker of reduced susceptibility (Supplemental Table 1). With the exception of passage 8, modest fold changes were observed, with shifts in IC_50_ values ranging from 5.4- to 6.5-fold compared to the wild-type SARS-CoV-2 stock virus. In lineage 1, the passage 8 virus displayed a >10-fold shift in IC_50_ (greater than highest concentration tested). Sequence analysis detected an identical 4 amino acid insertion in the N-terminal domain (215-216insKLRS) and 5 amino acid deletion in correspondence of the furin cleavage site (675-679del) in both lineages at all passages sequenced, as well as the amino acid substitution E340A in lineage 1, and R682W, and V1128F in lineage 2. The deletion at amino acids 675-679 has been previously described during passaging of SARS-CoV-2 in tissue culture suggesting enrichment to be a result of cell culture adaptation^43^ while the 215-216insKLRS was detected in the input virus. Neither 215-216insKLRS nor R682W variants were highly enriched with passaging (Supplemental Table 1) and enrichment of 675-679del and V1128F did not profoundly alter the VIR-7832 IC_50_. However, appearance of the E340A variant at 98.7% did correlate with a >10-fold shift in IC_50_ suggesting this variant may confer resistance.

To evaluate whether amino acid variants identified in the resistance selection conferred reduced susceptibility to sotrovimab and VIR-7832, neutralization of pseudotyped viruses encoding the S variants was assessed (Supplemental Table 2). Sotrovimab and VIR-7832 neutralized R682W and V1128F SARS-CoV-2 pseudotyped virus spike variants with IC_50_ values similar to wild type (< 2-fold change in IC_50_) indicating that these variants do not alter susceptibility. In contrast, E340A conferred reduced susceptibility to VIR-7831 and VIR-7832 (> 100-fold change in IC_50_) indicating that E340A is a VIR-7831/VIR-7832 monoclonal antibody resistance mutation (MARM).

As sotrovimab /VIR-7832 demonstrated a unique in vitro resistance profile, we investigated the potential for cross-resistance to MARMs that confer reduced susceptibility to the monoclonal antibodies bamlanivimab, imdevimab and casirivimab ^10, 11, 44–46^ using pseudotyped virus. Notably, some of these mutations are found in highly prevalent variants of concern^47–49^. Sotrovimab effectively neutralized pseudotyped viruses expressing spike MARMs that alter bamlanivimab, casirivimab and/or imdevimab activity (Table 3). Fold changes in IC_50_ values compared to wild-type were <3-fold for 18/19 variants tested. A modest 3.4-fold shift in the sotrovimab IC_50_ was observed for the V445A variant that confers reduced susceptibility to imdevimab. These data indicate that sotrovimab/VIR-7832 does not display cross-resistance with currently authorized mAbs and supports the potential combination use of sotrovimab/VIR-7832 with other mAb therapeutics.

**Table 3.** Storovimab and VIR-7832 retain activity against variants that confer resistance to authorized or approved mAbs. Activity of sotrovimab against variants conferring reduced susceptibility to bamlanivimab, imdevimab or casirivimab in a VSV/VeroE6 pseudotyped virus system. The geometric mean of IC_50_s and average fold-change versus the relative wild-type control from at least two independent experiments are shown.

| Amino Acid position | Substitution / Deletion | mAb with Reduced Susceptibility | Variants in Tested Spike Sequence | Sotrovimab IC<sub>50</sub> (ng/ml) | Average Fold Change in IC<sub>50</sub> Compared to Relative Wild-Type |
| --- | --- | --- | --- | --- | --- |
| E406 | W | casirivimab+<br>imdevimab | E406W | 53.66 | 0.74 |
| K417 | E | casirivimab | K417E | 67.71 | 0.89 |
| N439 | K | imdevimab | N439K,<br>D614G | 17.05 | 0.86 |
| N440 | D | imdevimab | N440D | 80.47 | 1.29 |
| N440 | K | imdevimab | N440K, D614G | 19.99 | 0.48 |
| K444 | Q | imdevimab | K444Q | 79.68 | 1.11 |
| V445 | A | imdevimab | V445A | 41.74 | 3.38 |
| G446 | V/I | imdevimab | G446V,<br>D614G | 18.41 | 1.50 |
| Y453 | F | casirivimab | G261D, Y453F | 27.28 | 2.19 |
| L455 | F | casirivimab | L455F, D614G | 21.65 | 0.56 |
| G476 | S | casirivimab | G476S | 36.97 | 2.94 |
| E484 | K | bamlanivimab | E484K, D614G | 12.91 | 0.33 |
| F486 | V/I | casirivimab | F486V | 82.24 | 1.10 |
| Y489 | H | casirivimab | Y489H | 92.29 | 1.48 |
| F490 | S | bamlanivimab | F490S | 33.10 | 0.85 |
| Q493 | K | casirivimab,<br>bamlanivimab | Q483K | 69.79 | 0.98 |
| S494 | P | casirivimab,<br>bamlanivimab | S494P, D614G | 29.10 | 2.50 |

### The sotrovimab/VIR-7832 epitope is highly conserved among SARS-CoV-2 sequences

The parental antibody of sotrovimab and VIR-7832 (S309) binds to a highly conserved sarbecovirus epitope that is potentially intolerant of variation. To investigate the current state of epitope conservation, >4,500,000 spike sequences from SARS-CoV-2 deposited in the GISAID database as of November 11, 2021 were examined for epitope variation. More than 99.6% conservation is seen for those amino acids comprising the epitope among currently available sequences for all positions including 15/23 amino acid positions that were ≥99.99 conserved (Table 4).

**Table 4.** The sotrovimab/VIR-7832 epitope is highly conserved. Conservation data comprising >4,500,000 sequences from the GISAID database and variants at each position are shown. Substitutions in green were tested in a pseudotyped virus assay.

| Amino Acid Position | Reference Amino Acid<sup>a</sup> | Substitutions Identified in order of Prevalence<sup>b</sup> | Percent Reference AA Conservation |
| --- | --- | --- | --- |
| 332 | I | V, T, N, D, S | >99.99 |
| 333 | T | I, K, A, S, P, R | >99.99 |
| 334 | N | K, Y, S, H, D, T, -, E, I | >99.99 |
| 335 | L | F, V, S, -, G, M | >99.99 |
| 336 | C | S*, -, P, R | >99.99 |
| 337 | P | L, S, T, H, R, -, K | >99.99 |
| 339 | G | D, S, C, V, F, -, N | 99.98 |
| 340 | E | D, K, A, G, Q, V, -, I | >99.99 |
| 341 | V | I, F, A, -, G, L, P, S | >99.99 |
| 343 | N | Y, -, D, K, S | >99.99 |
| 344 | A | S, T, V, D, F, G, -, P | 99.97 |
| 345 | T | S, I, -, N | >99.99 |
| 346 | R | K, S, I, G, T, -, F | 99.66 |
| 354 | N | K, D, S, T, H, Y, I, -, G, P | 99.95 |
| 356 | K | R, N, E, Q, M, T, -, G, W, Y, I, KR | 99.98 |
| 357 | R | K, I, -, T, G, S, C, L, RIS | 99.95 |
| 358 | I | V, L, F, -, T*, A, E* | >99.99 |
| 359 | S | N, T, G, R, I, -, C, F | 99.98 |
| 360 | N | Y, S, -, K, D, F, T, A, L | >99.99 |
| 361 | C | T*, -, G*, N*, F*, R*, S*, Y | >99.99 |
| 440 | N | K, S, Y, T, D, I, E, -, F, H, R | 99.79 |
| 441 | L | R, F, I, P, V, -, Y, H, M, S, E, N | >99.99 |
| 509 | R | -, K*, S*, T*, I*, H | >99.99 |

AA = amino acid <sup>a</sup>NCBI reference sequence: YP\_009724390.1 <sup>b</sup>Dash indicates a stop codon. Asterisks indicate the substitution could not be evaluated due to low expression of the spike protein containing this substitution

Sotrovimab activity against viral mutants carrying single substitutions in the epitope was assessed in pseudotyped virus assays. VIR-7831 effectively neutralized epitope variants at most amino acid positions tested (Supplemental Table 5). A moderate shift in activity was observed for the K356T variant (5.9-fold shift in IC_50_). Variants at two positions, E340 and P337, resulted in significant IC_50_ shifts indicating reduced susceptibility to VIR-7831. Moderate shifts in potency were observed for P337H, P337T and E340G substitutions (5.13-, 10.62- and 18.21-fold, respectively) while more significant shifts in potency were observed for P337L/R/K and E340A/K/Q (at least >50-fold). Notably, these variants are detected in a low number of sequences and do not have a pattern that suggest emergence in the GISAID database (296 and 223 variant counts out of >4,500,000 sequences for P337 and E340, respectively). This observation is consistent with the possibility that substitutions at these positions come with a fitness cost to the virus.

### Sotrovimab reduces weight loss, total viral load and infectious virus levels in a hamster model of SARS-CoV-2 infection

To evaluate the efficacy of sotrovimab in vivo, the hamster model was utilized. As it was unknown what effect the LS mutation would have in the hamster, a non-LS version of sotrovimab (SGHmAb-no-LS) was used for these experiments. Hamsters were administered SGHmAb-no-LS intraperitoneally at Day -1 (30, 5, 0.5 or 0.05 mg/kg) or Day -2 (15, 5, 0.5 or 0.05 mg/kg) prior to intranasal SARS-CoV-2 inoculation (Figure 3a). Using body weight as a marker of degree of clinical disease, doses of ≥5mg/kg resulted in significantly reduced weight loss at Day 4 compared to controls. (Figure 3b-e). Significant decreases in lung viral load were also observed at ≥5mg/kg as measured by RT-qPCR (Figures 3f-g). Day 4 TCID_50_ measurements indicated that antibody administered at ≥0.5 mg/kg resulted in significantly lower levels of infectious virus in lung tissue compared to controls (Figure 3h-i). Notably, across these experiments, no enhancement of disease was observed in animals receiving SGHmAb-no-LS based on changes in weight, viral RNA in the lungs, or TCID_50_ infectious virus levels. Collectively, these data indicate that sotrovimab prevented in a dose-dependent fashion virus replication and morbidity in SARS-CoV-2 challenged hamsters without signs of ADE at any dose tested.

**Figure 3.**
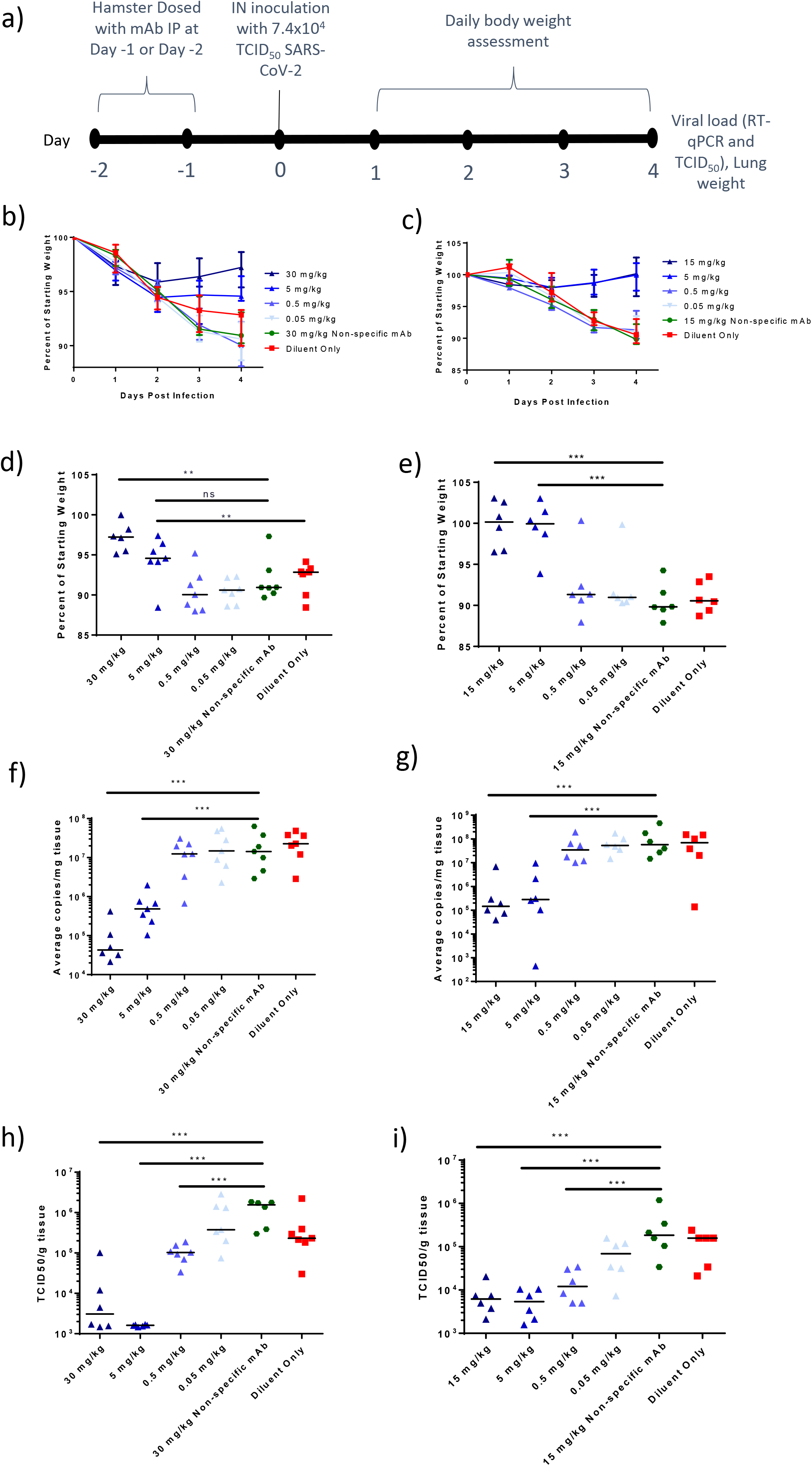
Sotrovimab (VIR-7831) shows in vivo efficacy in a hamster SARS-CoV-2 model of infection. a) Overview of hamster in vivo study design. b) and c) Animal weight over time as a percent of starting weight in animals dosed a Day -1 (b) or Day -2 (c). Medians of at least N=6 animals and interquartile range are shown. d) and e) Day 4 terminal weights expressed as a percentage of starting weight for animals dosed at Day -1 (d) or Day -2 (e). Bar denotes median values. f) and g) Day 4 lung viral load in Day -1 (f) or Day -2 (g) treated animals as assessed by RT-qPCR. Bar denotes median values. h) and i) infectious virus in lung at Day 4 for Day -1 (h) or Day -2 (i) dosed animals. Bar denotes median values. ns=not significant, ** = p<0.05, and *** = <0.005 as assessed by the Mann-Whitney U-test.

## DISCUSSION

Here we show the in vitro and in vivo preclinical characterization of sotrovimab and VIR-7832^50–52^. Both antibodies demonstrate high-affinity binding to S in vitro, including on the surface of cells, and effectively neutralize wildtype SARS-CoV-2 in a live virus assay. Sotrovimab and VIR-7832 neutralized all authentic virus pseudotyped virus variants tested in vitro. Sotrovimab neutralized 21 of 23 variants tested in a pseudotyped virus assay with a fold-change in IC_50_ of ≤3.3-fold, including the Omicron BA.1 and BA.1.1 sublineages. A shift in activity was observed for the Omicron BA.2 and BA.3 variants (16-fold and 7.3-fold change in IC_50_, respectively). Neutralization data from authentic virus assays were overall similar to those data generated from pseudotyped virus assays. Interestingly, higher shifts were observed in the sotrovimab IC_90_ versus the IC_50_ compared to the wild-type control (35.1-fold versus 15.7-fold, respectively) for the Omicron BA.2 sublineage. For other authentic virus variants, IC_50_ and IC_90_ changes compared to control were similar. This difference could be due to changes in assay conditions necessary for growth of Omicron BA.2 in vitro or could reflect a unique biology of this variant.

Preclinical studies indicate neutralization alone is unlikely to fully describe the activity of antibodies like sotrovimab, that retain effector function^17, 53^. Recent data demonstrate that despite a moderate reduction of activity observed for the sotrovimab parent molecule, S309, against the Omicron BA.2 variant in vitro, S309 displayed potent antiviral activity in a mouse model of Omicron BA.2 infection. This in vivo antiviral effect was abrogated when an effector function-deficient version of S309 was used, suggesting that assessment of neutralization activity alone does not capture the full antiviral capacity of antibodies with intact effector function like sotrovimab, and that Fc-mediated antiviral effects may also contribute to antibody efficacy. In addition, dose and pharmacokinetics are also important factors for determining clinical activity.

The sotrovimab/VIR-7832 epitope remains highly conserved among available sequences of circulating virus with ≥99.66% conservation of epitope amino acids. This is consistent with the value of the strategy used for isolation of monoclonal antibodies that neutralize both SARS-CoV and SARS-CoV-2 based on the idea that these two virulent human viruses are phylogenetically divergent within the sarbecovirus subgenus. Furthermore, MARMs identified at positions P337 and E340 are present at very low levels among current sequences. That amino acids P337 and E340 remain ≥99.99% conserved at this stage of the pandemic indicates that variants at these positions may confer disadvantageous effects on the virus, consistent with the conservation of this epitope across the sarbecovirus family^20^.

The vaccines presently being deployed around the world generate high-titer neutralizing antibodies that target the S protein RBM. Importantly, the RBM is highly immunodominant for responses to natural infection^55^. Vaccine-induced and convalescent immunity may therefore potentially put further mutational pressure on the RBM sequence to evade such antibody responses. In contrast, antibody responses overlapping with the sotrovimab/VIR-7832 epitope are limited after infection ^55^, possibly because of the shielding effect of the highly conserved N343 glycan. In this regard the epitope may face less vaccine-or infection-generated immune pressure, potentially preserving this conserved epitope long-term.

Recent data have indicated that the cells used to generate live virus stocks and overexpression of ACE2/TMPRSS2 in target cells used for assays can affect mAb activity in vitro^28, 56^. The sotrovimab/VIR-7832 parental antibody S309 seems particularly sensitive to in vitro methods using ACE2 overexpressing cells^56^. It is therefore notable that sotrovimab displays significant efficacy in an in vivo proof-of-concept SARS-CoV-2 infection experiment using hamsters despite the fact that patterns of engagement of hamster FcRs by human IgG1 antibodies may not reflect patterns of human IgG1 antibodies with their cognate human FcRs. These findings argue that in vitro data derived from such ACE2 overexpression cell lines do not accurately reflect the in vivo antiviral capacity of tested mAbs. Furthermore, that the significant in vivo effects of sotrovimab in the hamster model likely occurred in the absence of full effector functions due to species-specific interactions between antibodies and FcRs, argues that effects in COVID-19 patients incorporating both the neutralization capacity of the antibody plus the ability to harness the strength of the immune system could lead to positive clinical outcomes.

The clinical potential of VIR-7832, with the inclusion of the GAALIE Fc mutation, is of special interest in the context of SARS-CoV-2 infection. Previously published data by the Ravetch laboratory comparing the in vivo efficacy of a hemagglutinin-targeting mAb with and without inclusion of the GAALIE mutation in a transgenic humanized FcγR mouse model of influenza infection demonstrated superior efficacy of the GAALIE-containing antibody in both therapeutic and prophylactic experiments^25^. These effects were mediated by protective CD8^+^ T cell responses elicited by the GAALIE antibody. Clinical data examining the contribution of the adaptive immune response in SARS-CoV-2 infection indicate that poor T cell induction correlates with severe disease (reviewed in ^57^). Thus, the potential for VIR-7832 to augment the T cell response to SARS-CoV-2 infection could conceivably play a crucial role in limiting progression to severe COVID-19 disease or in treatment of severe established disease. This latter possibility is supported by recent publications showing that monoclonal antibodies with effector functions are especially effective in the therapeutic setting via recruitment of tissue-protective monocyte functions ^18^, and that potency of antibodies in the pre-clinical mouse model does not correlate with in vitro neutralizing activity of antibodies^17^.

Taken together, these data indicate that sotrovimab and VIR-7832 play a powerful role in the fight against COVID-19 through the dual action of broadly neutralizing activity paired with engagement of the immune system through effector function capabilities.

## METHODS

### Cells

Vero E6 cells (ATCC), Vero E6-TMPRSS2^58^ and Lenti-X 293T cells (Takara) were cultured in Dulbecco’s Modified Eagle’s medium (DMEM), 10% FBS, 1x Penicillin-Streptomycin at 37°C, 5% CO_2_.

### Monoclonal Antibodies

Sotrovimab (VIR-7831) and VIR-7832 were produced at WuXi Biologics (China). SGHmAb-no-LS, S309-LS, and S309-GRLR were produced at Humabs Biomed SA, a subsidiary of Vir Biotechnology (Bellinzona, Switzerland) in expiCHO cells transiently co-transfected with plasmids expressing the heavy and light chain, as previously described ^59^

### Virus

**S**ARS-CoV-2 isolates USA-WA1/2020, UK/VUI/3/2020, hCoV-19/South Africa/KRISP-K005325/2020, hCoV-19/Japan/TY7-503/2021, hCoV-19/USA/CA-SU-15-S02/2021 and hCoV-19/USA/PHC658/2021, hCoV-19/USA/MD-HP20874/2021, and hCoV-19/USA/MD-HP24556/2022 were obtained from BEI Resources. The isolate hCoV-19/Japan/UT-NCD1288-2N/2022 was obtained from the University of Wisconsin, Madison. To propagate SARS-CoV-2, VeroE6 or VeroE6-TMPRSS2 cells were seeded at 10X10^6^ cells in T175 flasks in growth media and infected the next day at a MOI of 0.001 in virus propagation media. Virus was adsorbed for 1 hour at 37°C. Virus inoculum was removed, flasks were washed once with PBS, 25 mL of infection media was added to the cells and flasks were incubated at 37°C. Supernatants were collected at 48 hours post-infection once cytopathic effect was visible, centrifuged at 500 x g for 5 minutes, followed by a second centrifugation at 1000 x g for 5 minutes. Clarified supernatants were then aliquoted and stored at -80°C. Virus titers were determined using a plaque assay on VeroE6 cells, using standard methods. Briefly, 10-fold dilutions of virus stock were incubated in 6 well plates with 2.4% colloidal cellulose overlay for 24 hours. Cells were fixed with 4% PFA for 30 minutes at room temperature (RT), permeabilized with 0.125% Triton X-100, stained with anti-SARS-CoV-2 nucleocapsid antibody at 1:5000 and goat anti-rabbit IgG HRP at 1:5000. Plaque forming units (PFU) were visualized with TrueBlue reagent.

### In vitro binding ELISA

For the ELISA assay, 96-well plates were coated with 100 µl/well recombinant SARS-CoV2 RBD diluted in assay diluent (1% BSA/PBS) at a final concentration of 2 μg/mL and incubated overnight at 4°C. Plates were washed three times with 300 µl/well wash buffer using an automated washer. Assay diluent (100 µl/well) was added to block the plates and incubated for 1 hour at room temperature (RT) with shaking. Assay diluent was removed, and plates washed three times with wash buffer. Serial 1:3 dilutions of mAb (concentration range from 6 µg/mL to 0.33 ng/mL) in assay diluent were dispensed at 100 µl/well and incubated 1 hour at RT with shaking, then washed three times with wash buffer. The HRP-conjugated secondary antibody reagent (1:5,000 dilution in assay diluent) was added to each well (100 µl/well) and incubated for 1 hour at RT with shaking. After three washes with wash buffer, 100 µl/well of 2-component TMB peroxidase substrate solution was dispensed in each well and developed for 5 minutes at RT. The reaction was stopped with 100 µL/well 1M H_2_SO_4_ and the OD was read immediately at 450 nm on a SpectraMax M5 Microplate reader. EC_50_ values were calculated using non-linear regression of log (agonist) versus response in Graph Pad Prism.

### Spike binding affinity quantification by SPR

Antibody was diluted to 2 µg/mL (1 mL) in HBS-EP+ buffer and injected at 10 µL/min for 30 seconds across one flow cell of a CM5 sensor chip immobilized with anti-human Fc antibody docked in a Biacore T200. SARS-CoV2-RBD diluted in HBS-EP+ buffer was then injected at a single concentration, 1:3 dilutions from 100 nM to 3.7 nM, across both the flow cell containing captured the antibody as well as a reference flow cell containing only anti-human Fc antibody. Binding was measured with a flow rate of 30 µL/min and an injection time of 600 seconds; dissociation was monitored for 1800 seconds after injection. Data were collected at 10 Hz. After each binding measurement, regeneration reagent was injected to prepare the surface for a new cycle. Experiments were performed at 25°C, with the samples held at 15 °C in the instrument prior to injection.

### Measurement of Binding to Human Fcγ Receptors by SPR

Binding of sotrovimab and VIR-7832 to human recombinant FcγRs was measured by surface plasmon resonance (SPR) on a Biacore T200. Briefly, Biotin CAPture Reagent (modified streptavidin) was injected across all flow cells of a CAP sensor chip docked in a Biacore T200. Biotinylated Fc receptors at 1 µg/mL were injected across a single flow cell at 10 µL/min for 60 seconds (one receptor per flow cell), with one flow cell reserved as a reference surface. VIR 7831 or VIR-7832 at 100 µg/mL (diluted in HBS-EP+) were injected across all flow cells for 200 seconds using a flow rate of 30 µL/min and association was monitored. Dissociation was monitored for another 200 seconds after injection. Data was collected at 10 Hz. After each binding measurement, CAP Regeneration reagent was injected to prepare the surface for a new cycle. Experiments were performed at 25°C, with the samples held at 15°C in the instrument prior to injection.

### Measurement of Binding to Human Complement Protein C1q

Binding of sotrovimab and VIR-7832 to human complement was measured by biolayer interferometry (BLI) using an Octet Red96 instrument (FortéBio). Briefly, anti-human Fab (CH1-specific) sensors were used to capture sotrovimab and VIR-7832 at 10 µg/ml for 10 minutes. The IgG-loaded sensors were then exposed to kinetics buffer containing 3 µg/ml of purified human C1q for 4 minutes, followed by a dissociation step in the same buffer for additional 4 minutes. Association and dissociation profiles were measured in real time as changes in the interference pattern.

### Binding to Cell Surface Expressed SARS-CoV-2 Spike Protein

The SARS-CoV-2 spike protein coding sequence (YP_009724390.1, Wuhan-Hu-1 strain) was cloned into a cell expression plasmid under the control of the human CMV promoter (phCMV1) to generate phCMV1 WT spike. ExpiCHO-S cells were seeded the day before transfection at 3 x 10^6^ cells/mL in ExpiCHO Expression Medium. Immediately before transfection, the cells were seeded at 6 x 10^6^ cells cells/mL in a volume of 15 mL in 125 mL shake flasks. Six μg of phCMV1 WT spike plasmid or vector control were diluted in 1.2 mL of iced OptiPRO SFM., followed by addition of 48 µL of ExpiFectamine CHO Reagent and complexing for 1 minute at RT. The transfection mixture was added dropwise to cells with gentle swirling. Cells were then incubated at 37°C, 8% CO2 with shaking for 42 hours. At 42 hours post-transfection, ExpiCHO-S cells were harvested, washed twice with FACS buffer and resuspended at a concentration of 1.0 x 10^6^ cell/mL in PBS. Cells (5 x 10^4^ cells in 50 µl/wells) were dispensed into a 96-well V-bottom plate. Antibody was serially diluted (1:4, 10 points) starting at a concentration of 10 µg/mL. Cells were pelleted at 300 x g for 5 minutes and resuspended in 50 µL/well of antibody serial dilutions and plates were incubated for 45 mins on ice. Cells were washed twice in FACS buffer. Alexa Fluor 647-labelled Goat Anti-Human IgG secondary Ab was diluted 1:750 in FACS buffer and 50 µL was added to the cell pellet for 15 min on ice. Cells were washed twice with FACS buffer, resuspended in 1% PFA. Data was acquired by flow cytometry (CytoFlex LX).

### Pseudotyped virus production

Lenti-X™ 293T cells were seeded in 10-cm dishes for 80% next day confluency. The next day, cells were transfected with the plasmid pcDNA3.1(+)-spike-D19 (encoding the SARS-CoV-2 spike protein) or pcDNA3.1(+)-spike-D19 variants using the transfection reagent TransIT-Lenti according to the manufacturer’s instructions. One day post-transfection, cells were infected with VSV-luc (rVSVΔG; Kerafast) at an MOI of 3. The cell supernatant containing SARS-CoV-2 pseudotyped virus was collected at day 2 post-transfection, centrifuged at 1000 x g for 5 minutes to remove cellular debris, aliquoted and frozen at -80°C.

### In Vitro Neutralization of SARS-CoV-2 Pseudotyped Virus

VeroE6 cells were seeded into flat bottom tissue culture 96-well plates at 20,000 cells/well and cultured overnight at 37°C. Twenty-four hours later, 9-point 1:4 serial dilutions of VIR-7831 were prepared in infection medium and each dilution was tested in triplicate per plate (range: 20,000 to 0.3 ng/mL final concentration). SARS-CoV-2 virus stock was diluted in infection media for a final concentration of 2000 plaque forming units per well (MOI 0.1). Antibody dilutions were added to virus and incubated for 30 minutes at 37°C. Media was removed from the VeroE6 cells, mAb-virus complexes were added, and cells were incubated at 37°C. At 6 hours post-infection, cells were fixed with 250 μL 4% PFA, incubated for 30 minutes at RT, then washed 3 times with PBS to remove residual PFA. The cells were permeabilized with 50 μL of 0.125% Triton X-100 in PBS for 30 minutes at RT. The blocking buffer was removed, 50 μL of SARS-CoV-2 nucleocapsid antibody at 1:2,000 in blocking buffer was added, and plate was incubated for 1 hour at RT. Plates were washed three times with PBS and then incubated for 1 hour at RT with 50 μL/well of goat anti-rabbit-Alexa647 secondary antibody at a final dilution of 1:1,000 mixed with 2 ug/mL Hoechst dye in blocking buffer. After washing 5 times with PBS, 100 μL of fresh PBS was added for imaging. Plates were imaged on a Cytation5 plate reader. Whole well images were acquired (12 images at 4X magnification per well) and nucleocapsid-positive cells were counted using the manufacturer’s software.

### Live virus neutralization

VeroE6 cells were seeded into flat bottom tissue culture 96-well plates at 20,000 cells/well and cultured overnight at 37°C. Twenty-four hours later, 9-point 1:4 serial dilutions of VIR-7831 were prepared in infection medium and each dilution was tested in triplicate per plate (range: 20,000 to 0.3 ng/mL final concentration). SARS-CoV-2 virus stock was diluted in infection media for a final concentration of 2000 plaque forming units per well (MOI 0.1). Antibody dilutions were added to virus and incubated for 30 minutes at 37°C. Media was removed from the VeroE6 cells, mAb-virus complexes were added, and cells were incubated at 37°C. At 6 hours post-infection, cells were fixed with 250 μL 4% PFA, incubated for 30 minutes at RT, then washed 3 times with PBS to remove residual PFA. The cells were permeabilized with 50 μL of 0.125% Triton X-100 in PBS for 30 minutes at RT. The blocking buffer was removed, 50 μL of SARS-CoV-2 nucleocapsid antibody at 1:2,000 in blocking buffer was added, and plate was incubated for 1 hour at RT. Plates were washed three times with PBS and then incubated for 1 hour at RT with 50 μL/well of goat anti-rabbit-Alexa647 secondary antibody at a final dilution of 1:1,000 mixed with 2 ug/mL Hoechst dye in blocking buffer. After washing 5 times with PBS, 100 μL of fresh PBS was added for imaging. Plates were imaged on a Cytation5 plate reader. Whole well images were acquired (12 images at 4X magnification per well) and nucleocapsid-positive cells were counted using the manufacturer’s software.

Modifications were made to the live virus neutralization assay to accommodate the differential growth kinetics of BA.2. VeroE6-TMPRSS2 cells were seeded into flat bottom tissue culture 96-well plates at 20,000 cells/well and cultured overnight at 37°C. Twenty-four hours later, 9-point 1:4 serial dilutions of VIR-7831 were prepared in FBS medium and each dilution was tested in triplicate per plate (range: 80,000 to 1.2 ng/mL final concentration). SARS-CoV-2 virus stock was diluted in BSA media for a final concentration of 200 plaque forming units per well (MOI 0.01). Antibody dilutions were diluted 1:10 into virus preparation and incubated for 30 minutes at 37°C. Media was removed from the cells, mAb-virus complexes were added, and cells were incubated at 37°C. At 18 hours (wild-type isolate) or 24 hours (Omicron variant isolates) post-infection, cells were fixed with 4% PFA, incubated for 30 minutes at RT, then washed 3 times with PBS to remove residual PFA. The cells were permeabilized with 100 μL of 0.25% Triton X 100 in PBS for 30 minutes at RT, followed by three washes with PBS. Cells were incubated with 50 µL of anti SARS-CoV-2 nucleocapsid antibody at 1:1000 in blocking buffer for 1 hour at RT. Plates were washed three times with PBS and then incubated for 1 hour at RT with 50 μL/well of goat anti-rabbit IgG Alexa647 secondary antibody at a final dilution of 1:1000 mixed with 2 µg/mL Hoechst dye in blocking buffer. After washing 3 times with PBS, 200 μL of fresh PBS was added for imaging. Plates were imaged on a Cytation5 plate reader. Whole well images were acquired (12 images at 4X magnification per well) and nucleocapsid-positive cells were counted using the manufacturer’s software. Two unique isolates of Omicron BA.2 derived from different sources were tested. Both isolates had identical spike sequences as determined by Sanger sequencing. Average IC_50_, IC_90_ and fold-change in activity values were determined from the combined data sets of both isolates.

### Determination of Viral Titer by Focus-Forming Assay

One day prior to infection, 1.2X10^4^ VeroE6 cells were plated in black-walled, clear bottomed 96-well plates. Virus samples were diluted 1:5 in infection media and adsorbed onto VeroE6 cells for one hour at 37°C. The cells were washed once and overlaid with 1% methylcellulose/serum-containing media. At 24 hours post-infection, the methylcellulose overlay was removed, and cells were washed with PBS. Cells were fixed with 4% PFA, incubated for 30 minutes at RT, then washed with PBS to remove residual PFA. The cells were permeabilized with 50 μL of 0.25% Triton X-100 in PBS for 30 minutes at RT. The Triton X-100 was removed, cells were washed twice with PBS, and incubated with 50 μL of SARS-CoV-2 nucleocapsid antibody at 1:2,000 in blocking buffer for one hour at RT. Plates were washed three times with PBS and then incubated for one hour at RT with 50 μL/well of goat anti-rabbit-Alexa647 secondary antibody at 1:1,000 in blocking buffer. After washing three times with PBS, 50 μL of Hoechst dye at 1:1,000 in PBS was added for imaging. Plates were imaged on a Cytation5 plate reader. Whole well images were acquired (12 images at 4X magnification per well) and nucleocapsid-positive foci were counted using the manufacturer’s software and used to determine focus-forming units/mL supernatant (FFU/mL).

### Determination of mAb-Dependent Activation of Human FcγRIIa, FcγRIIIa or FcγRIIb

Sotrovimab, VIR-7832, S309-LS, and a control mAb with abrogated FcγR binding, S309-GRLR, were serially diluted 6-fold in assay buffer from 10,000 ng/ml to 0.006 ng/ml. Nine-point serial dilutions of mAbs were incubated with 12,500 (for FcγRIIIa and FcγRIIb) or 10,000 (for FcγRIIa) CHO-CoV-2-Spike cells per 96-plate well in a white, flat-bottom plate for 15 minutes at room temperature. Jurkat effector cells expressing indicated FcγRs and stably transfected with an NFAT-driven luciferase gene were thawed, diluted in assay buffer, and added to the plate at an effector to target cell ratio of 6:1 for FcRγIIIa and FcγRIIb or 5:1 for FcγIIa. Control wells were also included that were used to measure antibody-independent activation (containing target cells and effector cells but no antibody) and background luminescence of the plate (wells containing assay buffer only). Plates were incubated for 18 hours at 37°C with 5% CO2. Activation of human FcγRs in this bioassay results in the NFAT-mediated expression of the luciferase reporter gene. Luminescence was measured with a luminometer after adding the Bio GloTM Luciferase Assay Reagent according to the manufacturer’s instructions. To control for background, the mean of the relative luminescence units (RLU) values in wells containing only Assay Buffer was calculated and subtracted from all data points. Data were expressed as the average of RLUs over the background

### Determination of NK-Cell Mediated Antibody-Dependent Cellular Cytotoxicity

Primary NK cell activation was tested using freshly isolated cells from two previously genotyped donors expressing homozygous low affinity (F158) or high affinity (V158) FcγRIIIa. Serial dilutions of mAbs (serially diluted 10-fold in AIM-V Medium from 40,000 ng/ml to 0.075 ng/ml) were incubated with 7,500 CHO-CoV-2 Spike cells per well of a 96 well round-bottom plate for 10 minutes. Target cell and antibody mixtures were then incubated with primary human NK cells as effectors at an effector-to-target ratio of 10:1. ADCC was measured using lactate dehydrogenase (LDH) release as a readout according to the manufacturer’s instructions (Cytotoxicity Detection Kit (LDH), Roche) after 4 hours of incubation at 37°C. In brief, plates were centrifuged for 4 minutes at 400 x g, and 35 µl of supernatant was transferred to a flat 384 well plate. LDH reagent was prepared and 35 µl were added to each well. Using a kinetic protocol, the absorbance at 490 nm and 650 nm was measured once every 2 minutes for 8 minutes, and the slope of the kinetics curve was used as result. The percent specific lysis was determined by applying the following formula: (specific release – spontaneous release) / (maximum release -spontaneous release) x 100.

### Determination of Monocyte-Mediated Antibody-Dependent Cellular Phagocytosis

ADCP assays were performed using human PBMCs freshly isolated from whole blood. CHO CoV-2-Spike cells were used as target cells and were fluorescently labeled with PKH67 Fluorescent Cell Linker Kit (Sigma Aldrich) prior to incubation with mAbs, according to manufacturer’s instructions. Serial dilutions of mAbs (serially diluted 5-fold from 5,000 ng/ml to 0.32 ng/ml in RPMI-1640 + L-glutamine supplemented with 10% Hyclone FBS + 2x anti-anti (antibiotic-antimycotic)) were incubated with 10,000 CHO-CoV-2-Spike cells per well of a 96 well polypropylene plate for 10 minutes. Primary PBMCs were fluorescently labeled with Cell Trace Violet according to the manufacturer’s instructions. Target cell and antibody mixtures were then incubated with labeled PBMCs at an effector-to-target ratio of 16:1. After an overnight incubation at 37°C, monocytes were stained with anti-human CD14-APC antibody (BD Pharmingen). Antibody-mediated phagocytosis was determined by flow cytometry, gating on CD14+ cells that were double positive for cell trace violet and PKH67. Raw data were exported from the flow cytometer into the flow cytometry analysis software FlowJo v10 (Becton Dickinson) for gating and determination of the percentage of CD14^+^ cells that were also double positive for cell trace violet and PKH67. Cells expressing only cell trace violet or only PKH67 were used to set the positive staining gates.

### In vitro resistance selection

The selection of variants in the presence of increasing concentrations of VIR-7832 was conducted in VeroE6 cells. The day before infection, 6 x 10^4^ VeroE6 cells were seeded in 24 well plates and incubated overnight at 37°C. The next day, 600 focus forming units (FFU) of SARS-CoV-2 virus (MOI = 0.01) was incubated with 0.5X IC_50_ of VIR-7832 (0.05 μg/mL) at 37°C for one hour in infection media. The mAb-virus complexes were adsorbed on VeroE6 cells for one hour at 37°C in duplicate wells. After adsorption, cells were washed with DMEM and overlaid with infection media containing 0.05 μg/mL VIR-7832. Control wells infected without antibody were included with each passage. Infected cells were monitored visually for CPE daily. In general, when infected cells exhibited ≥ 50% CPE, the culture supernatants were harvested, diluted 1:200, and added to fresh VeroE6 cells in 24-well plates with equivalent or increasing concentrations of VIR-7832. At each passage, supernatant was aliquoted and frozen at -80°C for titer and neutralization analyses.

### In vitro assessment of potential for ADE

VeroE6 cells were plated at 1.25X10^4^ cells/well one day prior to infection. For each independent experiment, moDCs and PBMCs from five unique moDC donors and six unique PBMC donors were used, with three unique donors used for each independent experiment. Cryopreserved monocytes from unique donors were differentiated into moDCs for six days using human moDC differentiation media according to the manufacturer’s protocol. Cryopreserved PBMCs from unique donors are thawed in the presence 0.3 mg/mL DNase and cultured in media for one day prior to infection. On the day of infection, moDCs, PBMCs, and U937 cells were counted and plated at 7.5X10^4^ cells/well.

To examine viral entry, 24 hours post-infection, cells were fixed with 4% PFA, incubated for 30 minutes at RT, then washed with PBS to remove residual PFA. The cells were permeabilized with 50 μL of 0.25% Triton X-100 in PBS for 30 minutes at RT. The Triton X-100 was removed, cells were washed twice with PBS, and incubated with 50 μL of SARS-CoV-2 nucleocapsid antibody at 1:2,000 in blocking buffer for one hour at RT. Plates were washed three times with PBS and then incubated for one hour at RT with 50 μL/well of goat anti-rabbit-Alexa647 secondary antibody at 1:1,000 in blocking buffer. After washing three times with PBS, 50 μL of Hoechst dye at 1:1,000 in PBS was added for imaging. Plates were imaged on a Cytation5 plate reader. Whole well images were acquired (12 images at 4X magnification per well) and nucleocapsid-positive cells were counted using the manufacturer’s software. The percent of nucleocapsid+ cells was quantified using the Gen5 Imager software (Biotek, Vermont) as number of Cy5+ cells, [(nucleocapsid+ cells)/number of Hoechst+ cells (total cells)]x100. Data was analyzed using Prism v8.00 (GraphPad Software, La Jolla California USA, www.graphpad.com).

In order to quantify chemokines and cytokines from supernatants in a BSL2 laboratory, supernatants were inactivated by 10 minutes exposure to UVC light at 5,000 µJ/cm2. Supernatants were diluted 1:5 in infection media and levels of cytokines/chemokines were quantified using the U-plex 96-well assay according to the manufacturer’s protocol (Meso Scale Diagnostics, Rockville, MD). Quantification of cytokines and chemokines were determined based on an 8-point standard curve in duplicate, provided by the manufacturer. Cytokine data was analyzed using the Discovery Workbench v4.0.13 software (Meso Scale Diagnostics). Data was graphed and statistical analyses were conducted using Prism software.

### Sequencing of SARS-CoV-2 Spike Gene

To isolate nucleic acid from the supernatant of viral passages, 120 μL of cell supernatant was added to 360 μL of Trizol and stored at -80°C for further analysis. Trizol collected samples from viral passages where a shift in neutralization > 2-fold relative to wild type was detected were subjected to RNA isolation using PureLink RNA Mini Kit with the incorporation of on-column PureLink DNase Treatment, following manufacturer’s instructions. Reverse transcription reactions were performed with 6 μL of purified RNA and oligoT primers using the NEB ProtoScript II First Strand cDNA Synthesis kit, according to manufacturer’s instructions. The resulting cDNA was used as a template for PCR amplification of the spike gene using the KapaBiosystems polymerase (KAPA HiFi HotStart ReadyMix) with primers 5’ aattatcttggcaaaccacg-3’ and 5’ tgaggcttgtatcggtatcg-3’. Amplification conditions included an initial 3 minutes at 95°C, followed by 28 cycles with 20 seconds at 98°C, 15 seconds at 62°C and 72°C for 2 minutes, with a final 4 minutes at 72°C. PCR products were purified using AMPure XP beads following manufacturer’s instructions. The size of the amplicon was confirmed by analyzing 2 μL of PCR products using the Agilent D5000 ScreenTape System. Products were quantified by analyzing 1 μL with the Quant-iT dsDNA High-Sensitivity Assay Kit. Twenty ng of purified PCR product was used as input for library construction using the NEBNext Ultra II FS DNA Library Prep kit following manufacturer’s instructions. DNA fragmentation was performed for 13 minutes. NEBNext Multiplex Oligos for Illumina Dual Index Primer Set 1 was used for library construction, with a total of 6 PCR cycles. Libraries size was determined using the Agilent D1000 ScreenTape System and quantified with the Quant iT dsDNA High-Sensitivity Assay Kit. Equal amounts of each library were pooled together for multiplexing and ‘Protocol A: Standard Normalization Method’ of the Illumina library preparation guide was used to prepare 8 pM final multiplexed libraries with 1% PhiX spike-in for sequencing. The MiSeq Reagent Kit v3 (600-cycle) was used for sequencing the libraries on the Illumina MiSeq platform, with 300 cycles for Read 1, 300 cycles for Read 2, 8 cycles for Index 1, and 8 cycles for Index 2.

### Bioinformatics Analysis of Conservation

Available genome sequences for SARS-CoV-2 were downloaded from Global Initiative on Sharing All Influenza Data (GISAID; https://www.gisaid.org/) on June 4, 2021. Bat and pangolin sequences were removed to yield human-only sequences. The spike open reading frame was localized by aligning the reference protein sequence (NCBI reference sequence: YP_009724390.1) to the genomic sequence of isolates with Exonerate v.2.4.0. Coding nucleotide sequences were translated in silico using seqkit v.0.12.0. Multiple sequence alignment was performed using MAFFT v.7.455. Variants were determined by comparison of aligned sequences to the reference sequence using the R v3.6.3/Bioconductor v.3.10 package Biostrings v.2.54.0.

### In vivo studies

Syrian golden hamster studies were conducted at Lovelace Biomedical (Albuquerque, NM). Twelve-to sixteen-week-old male hamsters were interperitoneally administered a non-LS version of VIR-7831 (SGHmAb-no-LS), control antibody or diluent Day -1 or Day -2 prior to virus challenge. Animals were inoculated intranasally at Day 0 with 7.4×10^4^ TCID_50_ with SARS-CoV-2 (isolate USA-WA1/2020). Animals were also weighed once daily in the morning beginning on study Day -10 and continuing until the end of the study. Following euthanasia, RT-qPCR was performed on lung homogenates using quantitative real-time PCR methods targeting the SARS-CoV-2 N gene and the median tissue culture infections dose (TCID_50_) was determined per Lovelace internal methodology.

## Author Contributions

Conceived studies: A.L.C, C.H-D., F.A.P, D.M., M.S., L.S., A.T., L.A.P., S.H., G.S., H.W.V., D.C., C.M.H. Designed studies and experiments: A.L.C, C.H.D., F.A.P, D.M., M.S., M.L.A., E. D., B.G., J.D., L.R., A.C., A.S., R.S., J.W., N.C., E.C., S.L., C.C., D.P., C.S., J.N., A.P., A.W., S. L, N. F., L.S., A.T., L.A.P., S.H., G.S., H.W.V., D.C., C.M.H. Performed experiments: D.M., M.S., M.L.A., B.G., J.D., E.D., A.S., L.R., H.T., J.D., S.S., D.P., C.S., J.N., S. L., N. F., B.S., S.B., J.W., J.Z., H.K., A.C., M.M-R., A.P., A.W., N.C., E.C. Analyzed and interpreted data: A.L.C., C.H-D., F.A.L., D.M., M.S., M.L.A., B.G., J.D., D.P., C.S., J.N., E.L., A.S., R.S., L.R., H.T., B.S., S.B., J.W., J.Z., H.K., A.C., M. M-R., N.C., E.C., S.L., A.W., C.C., L.S., A.T., S.H., G.S., H.W.G, D.C., C.M.H. Prepared the manuscript with input from all authors: A.L.C., G.S., A.T., L.P., D.C., H.W.G., C.M.H.

---

**Supplemental Figure 1.**
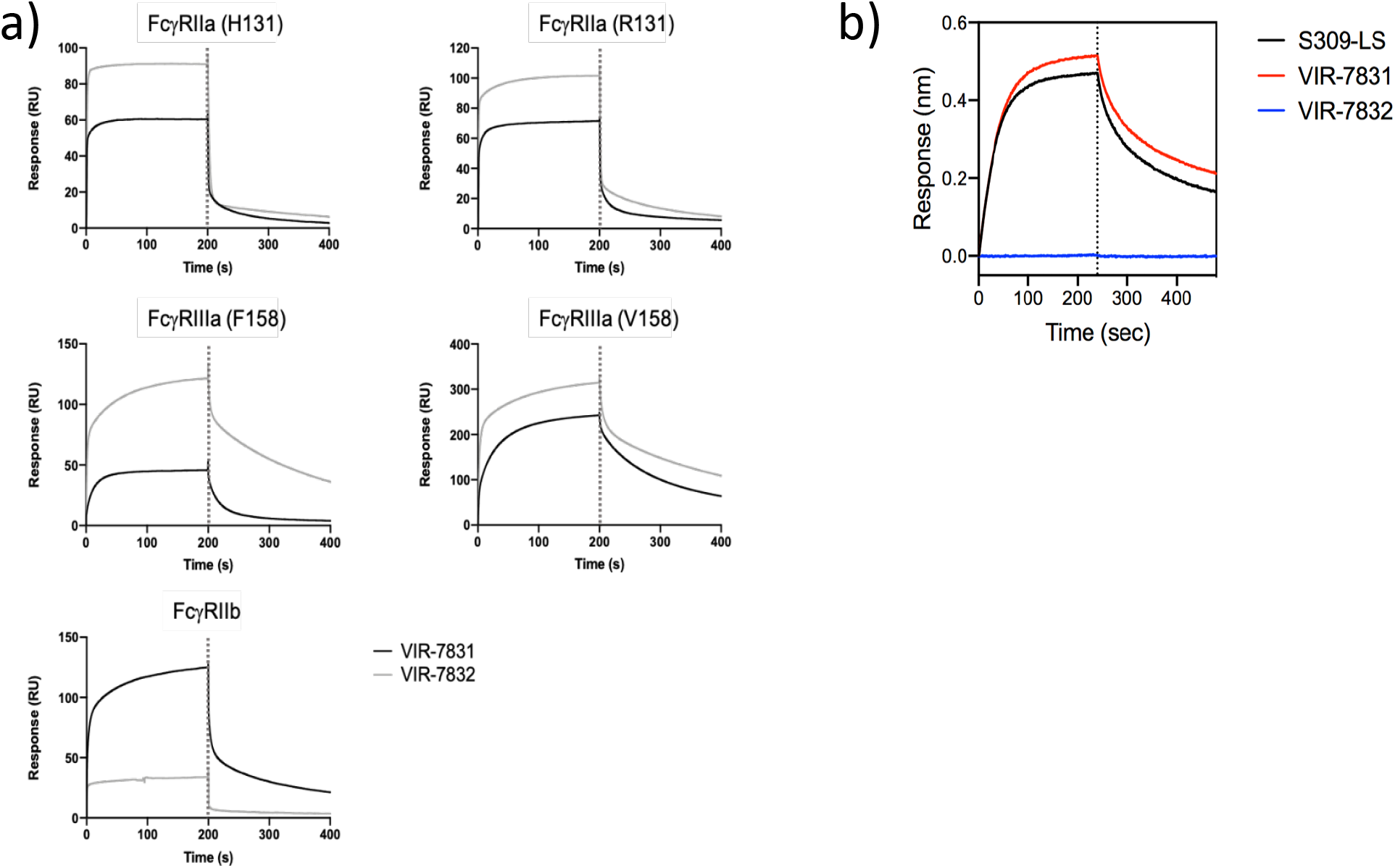
Binding of sotrovimab (VIR-7831) and VIR-7832 to human FcγRs and C1q as measured by SPR. Binding of sotrovimab (VIR-7831) and VIR-7832 to a) human FcγRIIa (H131 and R131 alleles), FcγRIIIa (F158 and V158 alleles) and FcγRIIb were measured using SPR. Biotinylated purified FcγRs were captured on the sensor chip surface prior to injection of sotrovimab (VIR-7831) or VIR-7832. Association and dissociation profiles (separated by the vertical dotted line) were measured in real time as change in the SPR signal. b) Binding of sotrovimab (VIR-7831) and VIR-7832 to complement component C1q was measured using BLI on an Octet Red96 instrument. Association and dissociation profiles (separated by the vertical dotted line) were measured in real time as change in the interference pattern.

**Supplemental Figure 2.**
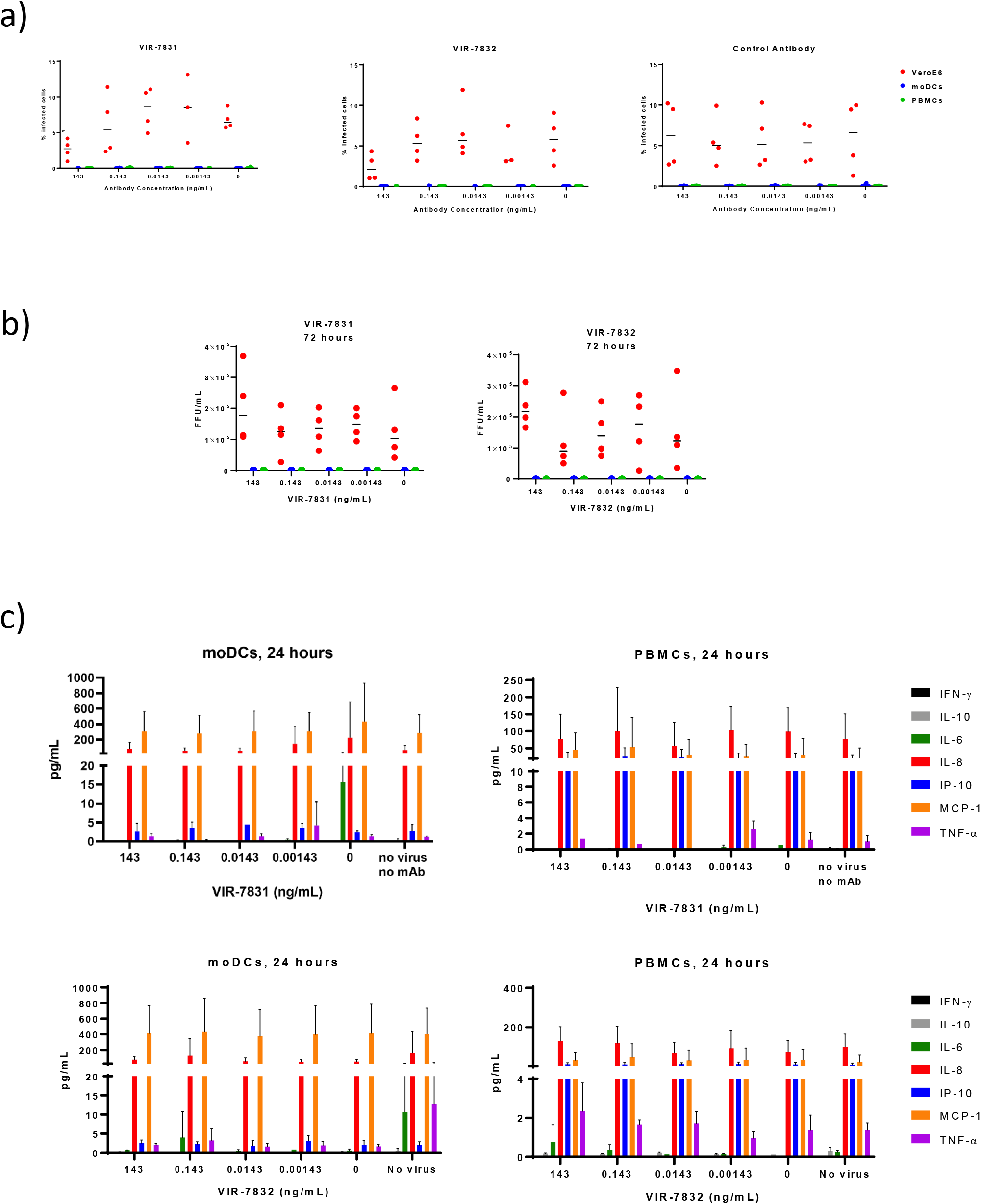
Sub-neutralizing concentrations of sotrovimab (VIR-7831) and VIR-7832 do not enhance viral entry, viral replication or cytokine production in vitro. Internalization (a) and replication (b) of SARS-CoV-2 was evaluated in VeroE6, moDCs or PBMCs at various timepoints. Two independent experiments with human moDCs and PBMCs from three individual donors were analyzed (5 unique moDC donors, 6 unique PBMC donors total between two experiments). VeroE6 cells were run in duplicate for both independent experiments. Data from each replicate well from two independent experiments are plotted as individual points, with horizontal lines representing the median. Mann-Whitney U-test comparison to no antibody group, *p<0.05. c) Supernatant cytokine and chemokine levels as measured by MSD at the indicated time post infection. Data from two independent experiments (three replicates each, five unique donors) are plotted as the mean and SD.

**Supplemental Figure 3.**
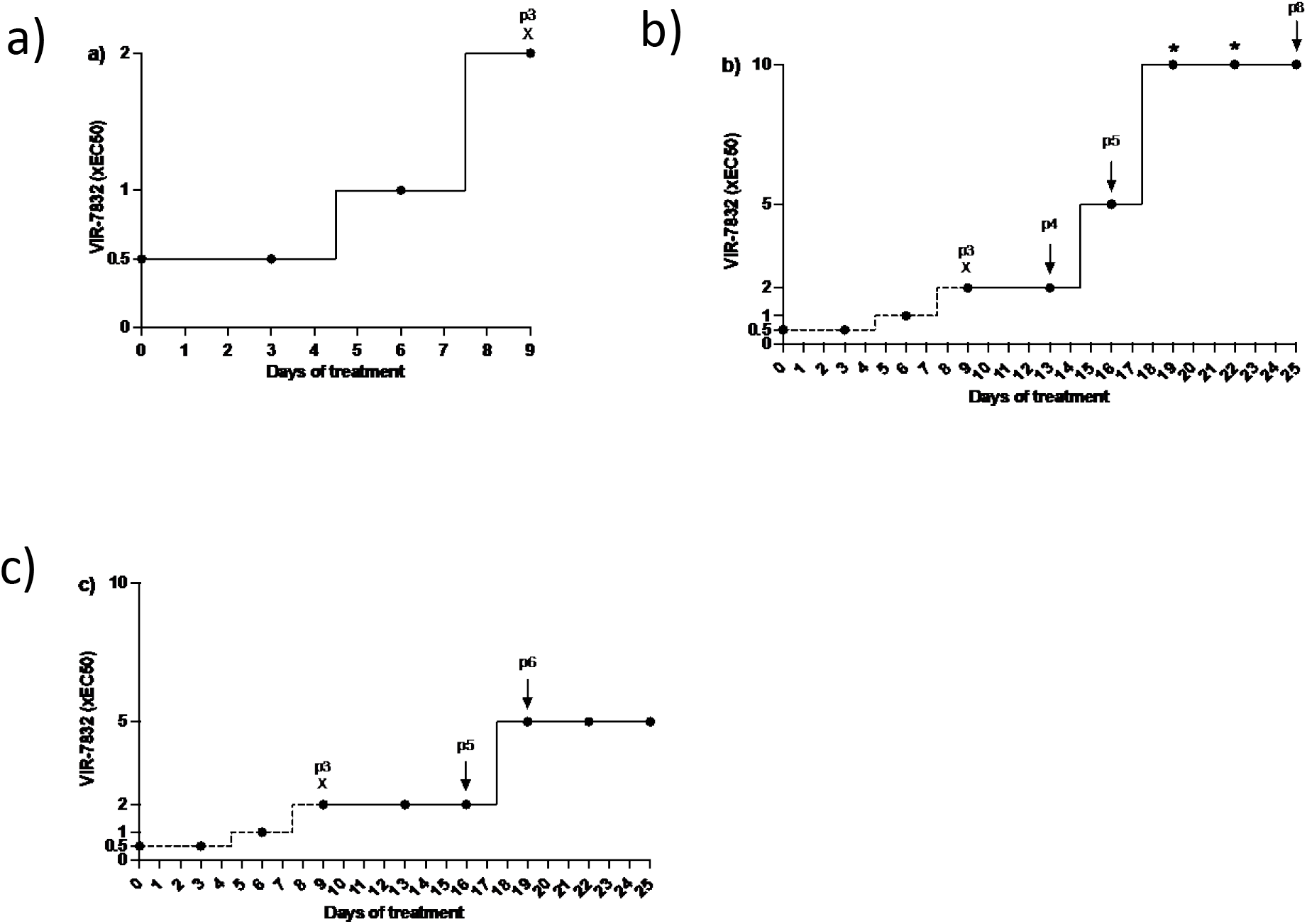
Overview of VIR-7832 resistance selection method. All passaging was conducted in duplicate wells. (a) VIR-7832 concentration was increased during each passage. P3 X indicates passage 3 virus, after which virus was lost with subsequent increases in concentration. In (b) and (c), p3X denotes where passage 3 virus from (a) was used to initiate (b) viral lineage 1 and (c) viral lineage 2. Arrows indicate passages that were subjected to sequence analysis, and * indicate the passages in lineage 1 with no detectable virus or CPE. Selection continued for a total of eight passages.

**Supplemental Table 1.** Amino acid substitutions identified in the SARS-CoV-2 S upon in vitro selection with VIR-7832. Spike gene sequences were compared to a SARS-CoV-2 reference sequence (NCBI: NC_045512.2) to identify variants. Fold-changes in IC_50_ were determined compared to the SARS-CoV-2 virus stock.

| Passage | Spike Gene Amino Acid Substitution (Freq)<sup>a,b</sup> | EC<sub>50</sub> (µg/mL) | Fold Change in EC<sub>50</sub> to WT<sup>c</sup> |
| --- | --- | --- | --- |
| SARS-CoV-2 virus stock<sup>c</sup> | H66R (5.7%)<br>N74K (12.6%)<br>T76I (5.6%)<br>215-216insKLRS (60.9%)<br>H655Y (3.1%) | 0.06 | NA |
| VIR-7832<br>Lineage 1, passage 4 | 215-216insKLRS (74.5%)<br>675-679 del (20.6%) | 0.34 | 5.64 |
| VIR-7832<br>Lineage 1, passage 5 | 215-216insKLRS (74.6%)<br>675-679del (66.0%) | 0.35 | 5.93 |
| VIR-7832<br>Lineage 1, passage 8 | 215-216insKLRS (74.7%)<br>E340A (98.7%)<br>675-679del (84.5%) | ND | >10 |
| VIR-7832<br>Lineage 2, passage 5 | 215-216insKLRS (73.9%)<br>675-679del (47.3%)<br>R682W (4.9%)<br>V1128F (3.5%) | 0.32 | 5.40 |
| VIR-7832<br>Lineage 2, passage 6 | 215-216insKLRS (75.3%)<br>675-679del (74.2%)<br>R682W (4.9%)<br>V1128F (30.9%) | 0.39 | 6.54 |

**Supplemental Table 2.**
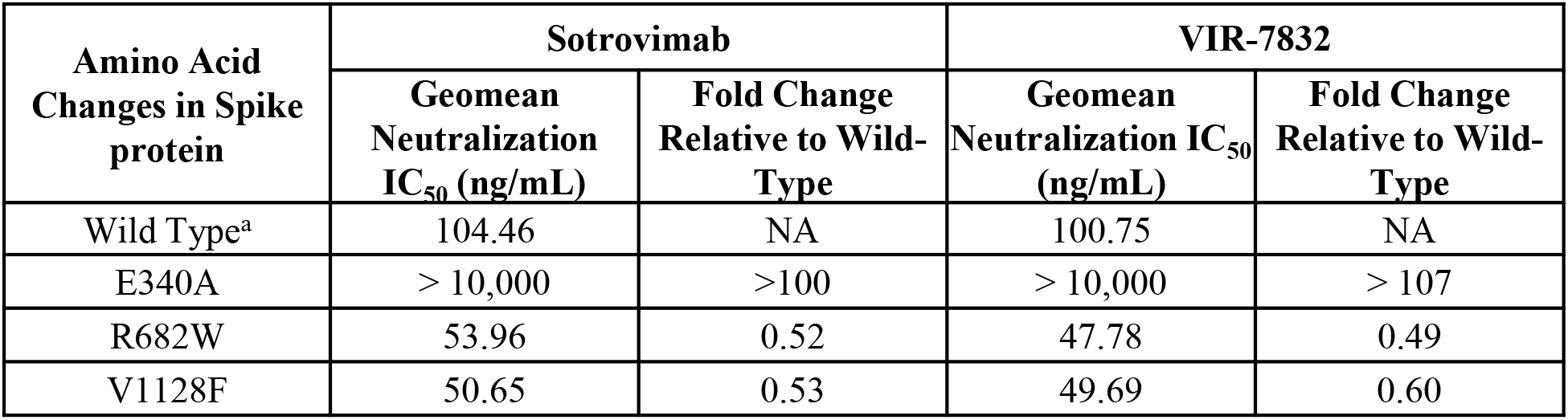
Sotrovimab and VIR-7832 activity against selected S variants. Sotrovimab/VIR-7832 epitope variants observed by in vitro resistance selection were individually tested in a VSV/VeroE6 pseudotyped virus system. The geometric mean of IC_50_s and average fold-change versus the relative wild-type control from at least two independent experiments are shown.

| Amino Acid<br>Changes in Spike<br>protein | Sotrovimab |  | VIR-7832 |  |
| --- | --- | --- | --- | --- |
|  | Geomean<br>Neutralization<br>IC<sub>50</sub> (ng/mL) | Fold Change<br>Relative to Wild-Type | Geomean<br>Neutralization IC<sub>50</sub><br>(ng/mL) | Fold Change<br>Relative to Wild-Type |
| Wild Type<sup>a</sup> | 104.46 | NA | 100.75 | NA |
| E340A | > 10,000 | >100 | > 10,000 | > 107 |
| R682W | 53.96 | 0.52 | 47.78 | 0.49 |
| V1128F | 50.65 | 0.53 | 49.69 | 0.60 |

**Supplemental Table 3.** Activity of sotrovimab against epitope variants. Sotrovimab/VIR-7832 epitope variants detected in sequences from the GISAID database were tested in a VSV/VeroE6 pseudotyped virus system. The geometric mean of IC_50_s and average fold-change versus the relative wild-type control from at least two independent experiments are shown.

| Epitope Reference Amino Acid | Amino Acid Substitutions in Spike Protein | Geomean Neutralization IC<sub>50</sub> (ng/mL) | Average Fold Change IC<sub>50</sub> Relative to Reference<sup>a</sup> |
| --- | --- | --- | --- |
| I332 | I332V, D614G | 53.43 | 1.48 |
|  | I332T, D614G | 66.74 | 1.61 |
| T333 | T333K, D614G | 22.14 | 0.67 |
|  | T333I, D614G | 30.93 | 0.59 |
| N334 | N334K, D614G | 45.36 | 1.27 |
|  | N334H, D614G | 30.74 | 0.87 |
|  | N334S, D614G | 34.37 | 1.13 |
|  | N334Y, D614G | 32.89 | 0.96 |
| L335 | L335F | 29.19 | 0.81 |
|  | L335S, D614G | 29.8 | 0.85 |
|  | L335V, D614G | 42.2 | 0.63 |
| P337 | P337H, D614G | 185.29 | 5.13 |
|  | P337L, D614G | >10000 | >192 |
|  | P337K, D614G | >10000 | >304 |
|  | P337R, D614G | >10000 | >192 |
|  | P337S, D614G | 127.69 | 1.26 |
|  | P337T, D614G | 383.27 | 10.62 |
| G339 | G339D, D614G | 117.38 | 1.18 |
|  | G339S, D614G | 32.67 | 0.86 |
|  | G339C, D614G | 68.79 | 1.18 |
|  | G339V, D614G | 78.32 | 1.26 |
| E340 | E340A | >10000 | >100 |
|  | E340D, D614G | 81.46 | 2.45 |
|  | E340K | >10000 | >297 |
|  | E340G, D614G | 640.07 | 18.21 |
|  | E340Q, D614G | >2500 | >50 |
|  | E340V, D614G | >10000 | >200 |
| V341 | V341I, D614G | 14.6 | 0.16 |
| N343 | N343S, D614G | 16.87 | 0.48 |

Supplemental Table 3 (con't)
| Epitope Reference Amino Acid | Amino Acid Substitutions in Spike Protein | Geomean Neutralization IC<sub>50</sub> (ng/mL) | Average Fold Change IC<sub>50</sub> Relative to Reference<sup>a</sup> |
| --- | --- | --- | --- |
| A344 | A344S | 92.19 | 0.89 |
|  | A344T, D614G | 40.94 | 0.62 |
|  | A344V, D614G | 53.87 | 1.14 |
|  | A344D, D614G | 131.53 | 2.57 |
| T345 | T345S, D614G | 42.92 | 0.69 |
|  | T345N, D614G | 20.29 | 0.57 |
|  | T345I, D614G | 11.78 | 0.20 |
| R346 | R346I, D614G | 65.39 | 1.72 |
|  | R346S, D614G | 42.87 | 1.13 |
|  | R346T, D614G | 89.04 | 1.25 |
|  | R346K, D614G | 24.76 | 0.72 |
|  | R346G, D614G | 43.73 | 0.85 |
| N354 | N354D | 104.8 | 1.00 |
|  | N354K, T95I | 70.62 | 0.76 |
|  | N354H, D614G | 74.88 | 1.06 |
|  | N354K, T95I | 70.62 | 0.76 |
|  | N354S, D614G | 61.78 | 0.89 |
|  | N354I, D614G | 37.09 | 1.05 |
|  | N354T, D614G | 43.32 | 0.65 |
|  | N354Y, D614G | 67.63 | 1.08 |
| K356 | K356A | 111.18 | 1.06 |
|  | K356R, D614G | 29.61 | 0.78 |
|  | K356N, D614G | 53.22 | 1.12 |
|  | K356M, D614G | 64.09 | 1.03 |
|  | K356T, D614G | 281.13 | 5.9 |
|  | K356E, D614G | 76.54 | 1.65 |
|  | K356Q, D614G | 43.2 | 0.94 |
| R357 | R357K, D614G | 39.13 | 1.02 |
|  | R357I, D614G | 46.41 | 0.98 |

Supplemental Table 3 (con't)
| Epitope Reference Amino Acid | Amino Acid Substitutions in Spike Protein | Geomean Neutralization IC<sub>50</sub> (ng/mL) | Average Fold Change IC<sub>50</sub> Relative to Reference<sup>a</sup> |
| --- | --- | --- | --- |
| I358 | I358V, D614G | 32.71 | 0.7 |
|  | I358L | 44.01 | 0.8 |
|  | I358F, D614G | 42.69 | 1.19 |
| S359 | S359N | 95.55 | 0.96 |
|  | S359G, D614G | 37.56 | 0.8 |
|  | S359R, D614G | 62 | 0.89 |
|  | S359T, D614G | 39.98 | 0.84 |
|  | S359I, D614G | 38.01 | 0.84 |
| N360 | N360S, D614G | 34.08 | 0.72 |
|  | N360Y, D614G | 29.67 | 0.61 |
| N440 | N440D | 80.47 | 1.29 |
|  | N440Y, D614G | 42.65 | 0.68 |
|  | N440S, D614G | 51.34 | 0.73 |
|  | N440K, D614G | 19.99 | 0.48 |
|  | N440I, D614G | 40.21 | 1.22 |
|  | N440T, D614G | 50.81 | 0.82 |
|  | N440E, D614G | 46.97 | 0.80 |
|  | N440F, D614G | 43.07 | 0.85 |
| L441 | L441F, D614G | 25.21 | 0.4 |
|  | L441I, D614G | 32.96 | 0.7 |
|  | L441R, D614G | 42.3 | 0.87 |
|  | L441V, D614G | 48.24 | 0.98 |
|  | L441P, D614G | 44.95 | 0.76 |

